# Persistent chromatin alterations and gene expression reprogramming follow widespread DNA damage in glioblastoma

**DOI:** 10.1101/2025.06.18.660431

**Authors:** Aram S. Modrek, Ken Chandradoss, Catherine Do, Ravesanker Ezhilarasan, Theodore Sakellaropoulos, Jerome Karp, Yingwen Ding, Ze-Yan Zhang, Melanie Graciani, Giulia Cova, Jennifer E. Phillips-Cremins, Erik P. Sulman, Jane A. Skok

## Abstract

DNA damage from routine cellular processes or exogenous insults can have a lasting impact on gene regulation beyond genetic mutations. The prevailing paradigm for the consequences of DNA damage repair revolves around restoration of the original genetic sequence, but long-term changes in chromatin configuration, gene expression and DNA modifications have not been analyzed. We introduced numerous, simultaneous Cas9-mediated DNA double strand breaks (DSBs) at defined locations in human glioblastoma cells and tracked both non-genetic and genetic alterations over time. Megabase-scale genomic alterations that endured two weeks after the initial damage were detected, involving a shift from transiently increased intra-TAD interactions to persistent long range *cis* and *trans* contacts, alterations in gene-expression and associated large structural variations. These findings reveal that widespread DNA damage, such as chemotherapy or radiotherapy, can trigger long-term genetic and non-genetic modifications which alter cellular function and may impact tumor outcome and the emergence of resistant cells.

## Main Text

Following treatment with DNA damaging therapy, tumors recur with new phenotypic traits that are not fully explained by genetic alterations or clonal selection alone (*1–3*). This discrepancy underscores the hypothesis that other mechanisms—such as persistent changes in DNA modifications, chromatin configuration, and gene expression—may play a role in fueling therapy resistance. Indeed, the DNA damage response mobilizes chromatin modifiers (*4, 5*), but whether these mediate changes that culminate in long-term gene expression shifts remains poorly understood.

DNA damage arises during physiological cellular processes and from exogenous sources such as chemotherapy, and therapeutic or environmental radiation. Glioblastoma presents a particularly compelling model to investigate DNA damage–induced changes as standard therapy includes 60 Gray of radiation, concurrent with use of the chemotherapeutic agent temozolomide (TMZ) and DNA damage generating Tumor Treating Fields (*6, 7*). The radiation portion alone is estimated to make a total of ~1,500 double-strand breaks per each cell (*8*). Despite multi-modal DNA-damaging therapy, tumors exhibit few new genetic lesions in driver oncogenes or tumor-suppressors, but recur with widespread, seemingly arbitrary, non-genetic alterations, as highlighted in recent clinical and preclinical work (*2, 9–11*). Therefore, elucidating mechanisms underlying damage-induced phenotypes could illuminate new strategies to prevent or counteract the emergence of resistant glioblastoma subclones following therapy.

Previous work has shown that chromatin and DNA modifications can change acutely upon radiation or endonuclease-induced breaks, but these studies were performed over short time frames (hours) and thus it is not known whether there is a long-term impact on chromatin and its 3D organization that leads to sustained changes in gene regulation (*4, 12–14*). Another limitation of these studies is the approach used for introducing DNA damage. For example, radiotherapy or chemotherapy can induce a variety of randomly localized DNA damage insults at different timepoints in individual cells, which prevents a clean analysis of longitudinal changes. Furthermore, use of fixed target homing endonucleases systems, which are unable to target methylated DNA likely produce a target selection bias, limiting the generalization and comprehensiveness of the mechanisms underlying the long-term effects of DSBs.

In order to dissect the dynamics of genetic and non-genetic events that follow DNA damage, it is important to study tumor evolution over controlled time periods without the confounding effects of the varied nature of chemoradiotherapy-induced breaks. The latter are randomly introduced into the genome in individual cells and cannot be tracked over time as part of bulk cell analysis. To address these limitations, we used a “multi-target” CRISPR–Cas9 approach that enables widespread pre-mapped DNA breaks at hundreds to thousands of loci in human glioblastoma stem cells (*15*). This approach, in similar fashion to prior efforts (*16, 17*), introduces DSBs at pre-defined sites (while mimicking the widespread nature of chemoradiotherapy-induced DNA damage) allowing us to systematically track the downstream effects on the genome, with the goal of studying the mechanisms underlying short and long-term changes in gene regulation. Following the introduction of DNA breaks, we examined the interplay between changes in gene expression in the context of DNA and chromatin modifications, transient and persistent changes in 3D chromatin folding and the emergence of large-scale structural variations.

## Results

### Genome-wide DNA damage leads to dynamic alterations in genome-organization and gene-expression over time

To gain insight into how cancer genomes could be altered following DNA-damaging therapies such as chemotherapy or radiation, we performed time course experiments following the simultaneous induction of multi-targeted DSBs. Glioma-Stem Cell lines (GSCs) (*18*) were co-infected with wt-spCas9 and two distinct doxycycline-inducible multi-target guide-RNAs (mtgRNA), termed mtg142 and mtg483, to simultaneously introduce numerous DSBs across the genome **(fig. S1A)**. These guides were selected because they have 142 and 483 predicted DSB target sites, respectively, which allows us to assess similar levels of damage to that incurred with cancer directed therapy. Details of mtgRNA design and generation are previously described (*15*). Induction of the mtgRNAs led to the formation of γH2AX foci as assessed by immunofluorescence at three hours, with each guide exhibiting similar toxicity to 2-4 Gy of radiotherapy as measured by colony formation assays in human glioma U87 cell lines **(fig. S1B and C)**. To create a double-strand break (DSB) map for back-tracing of genomic changes with damaged sites, we performed in-suspension break labeling in situ and sequencing (isBLISS) **(Fig. 1A and fig. S1D)**, which enabled us to generate a map of DSB sites for mtg142 and mtg483, which had 339 and 1,386 total verified cut sites, respectively, shown alongside predicted DSB target sites **(Fig. 1B)**. The increase of actual over predicted DSB sites occurs because of the presence of lower affinity sites with nucleotide mismatches.

**Figure 1.**
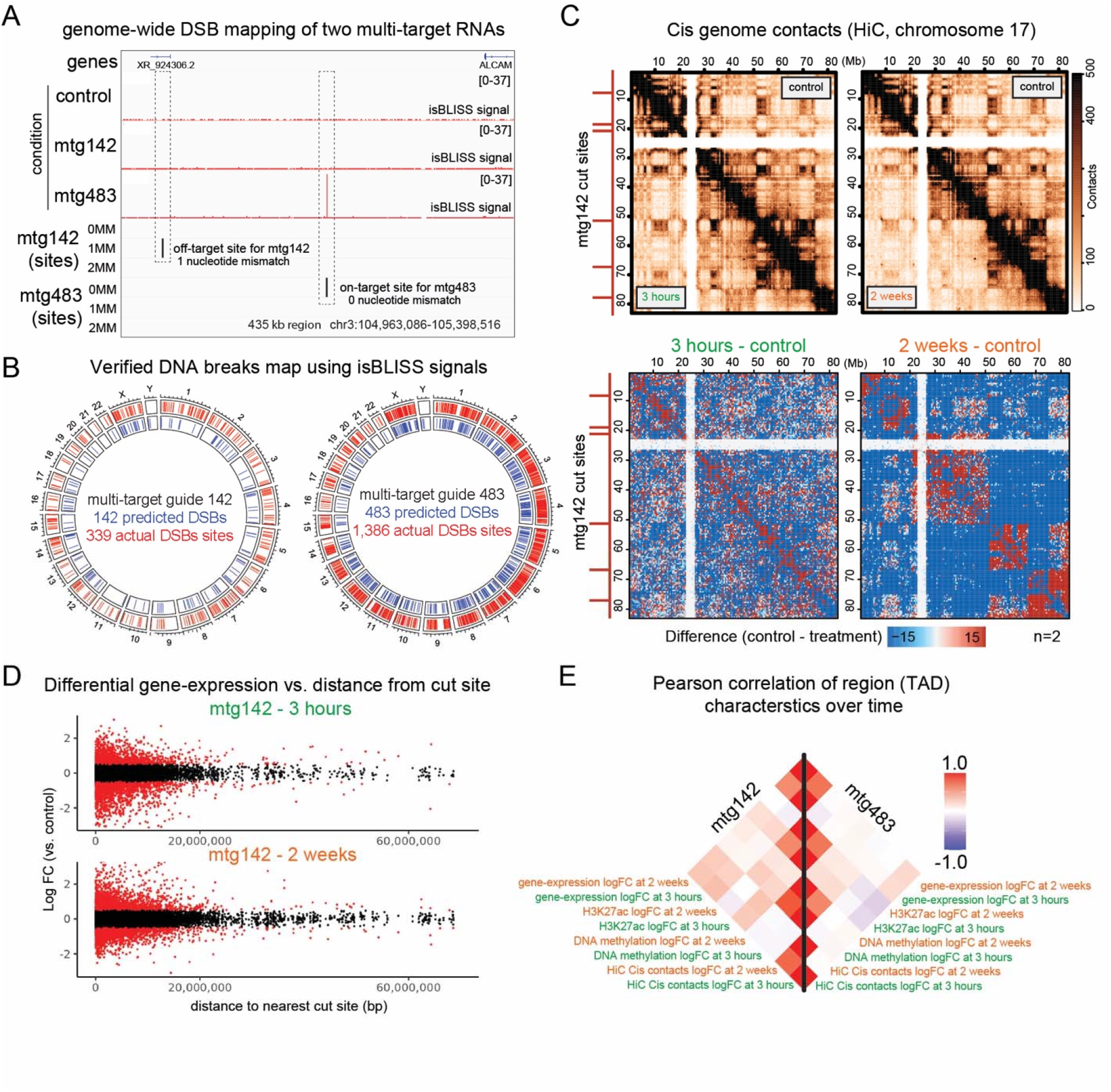
Genome-wide DNA damage leads to distal and dynamic alterations in genome-organization and gene-expression over time. A. Double-strand break (DSB) sequencing (isBLISS) of control (Cas9 only), CRISPR-Cas9 guides mtg142 and mtg483. Bottom rows (sites) represent regions that have either 0, 1 or 2 mismatches (MM) in nucleotides (theoretical or possible cut sites due to lower affinity off-targeting). B. Genome-wide DSB maps with isBLISS verified sites marked in red. Predicted sites (blue) represent perfect guide to genome matches. C. Hi-C (genome-contact) maps of chromosome 17 showing contacts (top panel) and contact changes (bottom panel) over time (3 hours and 2 weeks). Cut sites for mtg142 for this region are shown on the left y-axis in red. D. RNA-seq of differentially expressed genes marked as a function of distance to the nearest cut site for mtg142. E. Pearson correlation plot of aggregated log-fold changes in gene-expression, H3K27ac ChIP-seq, DNA methylation and Hi-C within Topologically Associated Domains (TADs).

We performed genome-wide Chromosome Conformation Capture with Hi-C at each of the three time points (control, 3 hours, and 2 weeks) to examine the longitudinal impact of DSBs generated by the two mtgRNAs on chromatin organization. Changes in *cis* genomic contacts were observed Megabases (Mb) from verified cut sites that evolved over a period of two weeks, suggesting a persistent re-programming event **(Fig. 1C and fig. S2A)**. Although the two guides target different sequences, the 3-hour and 2-week samples exhibited similar overall variance in gene expression levels as shown by principal component analysis **(fig. S2B**). Significant gene expression alterations were found at promoters of genes proximal to cut sites and up to 20 megabases away at the 3 hour time point and changes in mRNA levels persisted 2 weeks after the introduction of the DSBs (**Fig. 1D and fig. S2C**). As expected, gene-set enrichment analysis of the RNA-seq data highlight DNA-damage repair processes, but at both timepoints they also reveal processes involved in chromatin and DNA modifications **(fig. S3A to D and data S1 to S4)**. These findings suggest that the cell could be acutely and chronically upregulating chromatin modifying factors in response to DNA damage.

To determine the impact of DNA and chromatin modifying processes, we performed genome wide DNA methylation profiling and ChIP-seq for H3K27ac (**Fig 1E and fig. S4A to C)**. Pearson correlations showed that alterations in the signals of these data sets, which were averaged over topologically associated domains (TADs), could be correlated or anti-correlated between the two different time points and mtgRNAs used **(Fig. 1E)**. These findings demonstrate that the introduction of DSBs leads to context specific rewiring of the genome at the level of 3D interactions, chromatin/DNA modifications and gene expression and that reprogramming persists for 2 weeks. Furthermore, the results highlight that changes detected at 3 hours can be distinct from the longer-term alterations found at 2 weeks.

### Transient increases in cis chromatin contacts are followed by a delayed increase in trans chromatin interactions and compartment A-B intermingling

To investigate the connections between changes in 3D chromatin folding chromatin and chromatin activity, we used the H3K27ac and HiC data to determine whether shifting between transcriptionally active (A) and inactive (B) compartments occurred after the introduction of DSBs **(fig. S5A to C)**. Despite differences in the number and location of targeted DSBs, similar broad mega-base sized changes were detected within the regions near cut sites in cells expressing the two different guides **(Fig. 2A)**. A/B compartment traces were largely unchanged in placement between control and DSB-induced samples. However, it is noteworthy that the strength of the compartment score was attenuated upon the induction of DSBs, suggesting the dissolution of certain B compartments at some genomic locations. As shown in the example for chromosome 17, these were mostly positive compartment shifts, associated with mega-base scale alterations in chromatin contacts, **(Fig. 2B and fig. S5D)**. Compartment shifts can be associated with changes in regulatory marks, as illustrated in **Fig. 2C**, which shows an overall increase in H3K27ac, DNA methylation, and sustained gene-expression changes. Although major switches between compartments were not observed, a significant portion of compartments had score shifts that were widespread across the genome **(Fig. 2D)**. This change was significantly observed when comparing compartments with or without cut sites in the mtg142 condition at 2 weeks (p-value= 0.01, Fisher’s exact test), but not for mtg483. Compartments with shifts were also more prone to have alterations in chromatin contacts, DNA methylation and gene-expression, in contrast to unchanged compartments (compartments without a score shift at any timepoint) **(fig. S5F)**.

**Figure 2.**
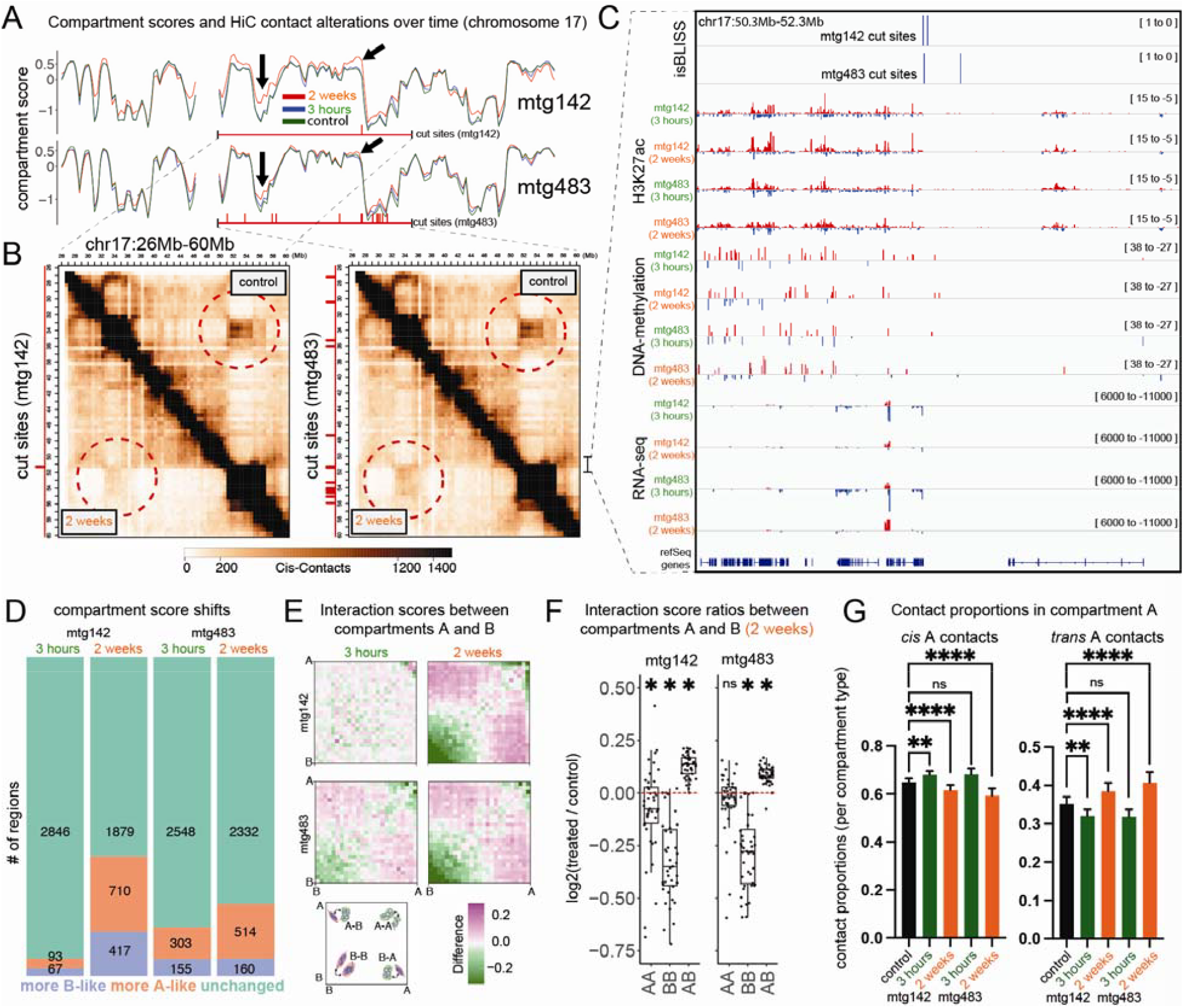
Transient increases in cis chromatin contacts are followed by a delayed increase in trans chromatin interactions and compartment A-B intermingling. A. Compartment scores over chromosome 17 for mtg142 and mtg483. Arrows point to areas of the genome that have a positive shift in compartment score. B. Contact maps (HiC) (2 weeks vs. control) show similar patterns of alterations. Red circles highlight areas of the genome that have significant changes in long-range cis contacts near areas of cut sites. C. Double-strand breaks induced by two different mtgRNAs show an increase in surrounding H3K27ac, DNA hypermethylation and longitudinal alterations in gene-expression in a 2 Mb region on chromosome 17. D. Distribution of compartment changes (absolute change >0.2 in score) over time and condition based on the number of genomic regions defined by 1 Mb windows. E. Saddle plots representing interaction scores between A and B compartments across the genome. F. Quantification of A/B interaction scores (represented as ratios) show an increase in A-B interactions (two-tailed one-spample t-test, * p-value < 0.01, ns = not significant) G. Contact proportions binned into *cis* or *trans* found in A compartments (ANOVA and Dunnett’s test, ** p-value < 0.01, **** p-value < 0.0001, ns = not significant).

These results imply that the introduction of double-strand breaks induces sustained dissolution of A and B compartments—where regions originally maintained in separate domains begin to intermingle. Importantly, in cells expressing both guide RNAs, saddle plots show that changes in A-B compartment interactions show subtle differences at 3 hours, but by 2 weeks reveal significant increases, indicating an increase in A-B compartment intermingling that reflects changes in DNA and chromatin modifications and alterations in gene expression (**Fig. 2E, F and fig. S6A, B**). To investigate further, we analyzed changes in *cis* and *trans* contacts. The proportion of *cis* contacts was transiently increased at 3 hours and subsequently decreased at 2 weeks, while the inverse trend was observed for trans-contacts, which exhibited a higher final proportion of interactions in both A and B compartments **(Fig 2G and fig. S6C)**.

In summary, induction of multiple double-strand breaks leads to transient increases in *cis* chromatin contacts associated with minimal compartment mixing, early sustained mega-base scale alterations in DNA methylation, H3K27 acetylation and gene expression **fig. S6D**. Persistent changes are accompanied by a delayed increase in *trans* chromatin interactions and a more prominent shift in compartment A-B intermingling.

### Reprogramming of long-range 3D genome contacts and regulatory elements following DNA damage over time are associated with persistent alterations in gene-expression

To investigate the connection between changes in chromatin folding and gene expression at granular resolution, we first analyzed intra- and inter-TAD contacts over time. As expected, control cells exhibited more frequent intra-TAD than inter-TAD interactions, consistent with the insulating properties of TADs—a configuration that persisted shortly after DNA damage. However, by 2 weeks post-induction, we observed a marked and reproducible increase in in long-range inter-TAD contacts spanning 10–20 Mb in both mtgRNA conditions (**Fig. 3A and fig. S7A**,**B**). Notably, this increase was only seen in cells that had DNA breaks, indicating that DSBs themselves drive these long-range contacts. Aggregate TAD analysis further revealed a transient increase in intra-TAD contacts at 3 hours (consistent with prior observations by the LeGube Lab), followed by a consistent decrease at 2 weeks (**fig. S7C**), suggesting a temporal shift from local to more extensive chromatin reorganization.

**Figure 3.**
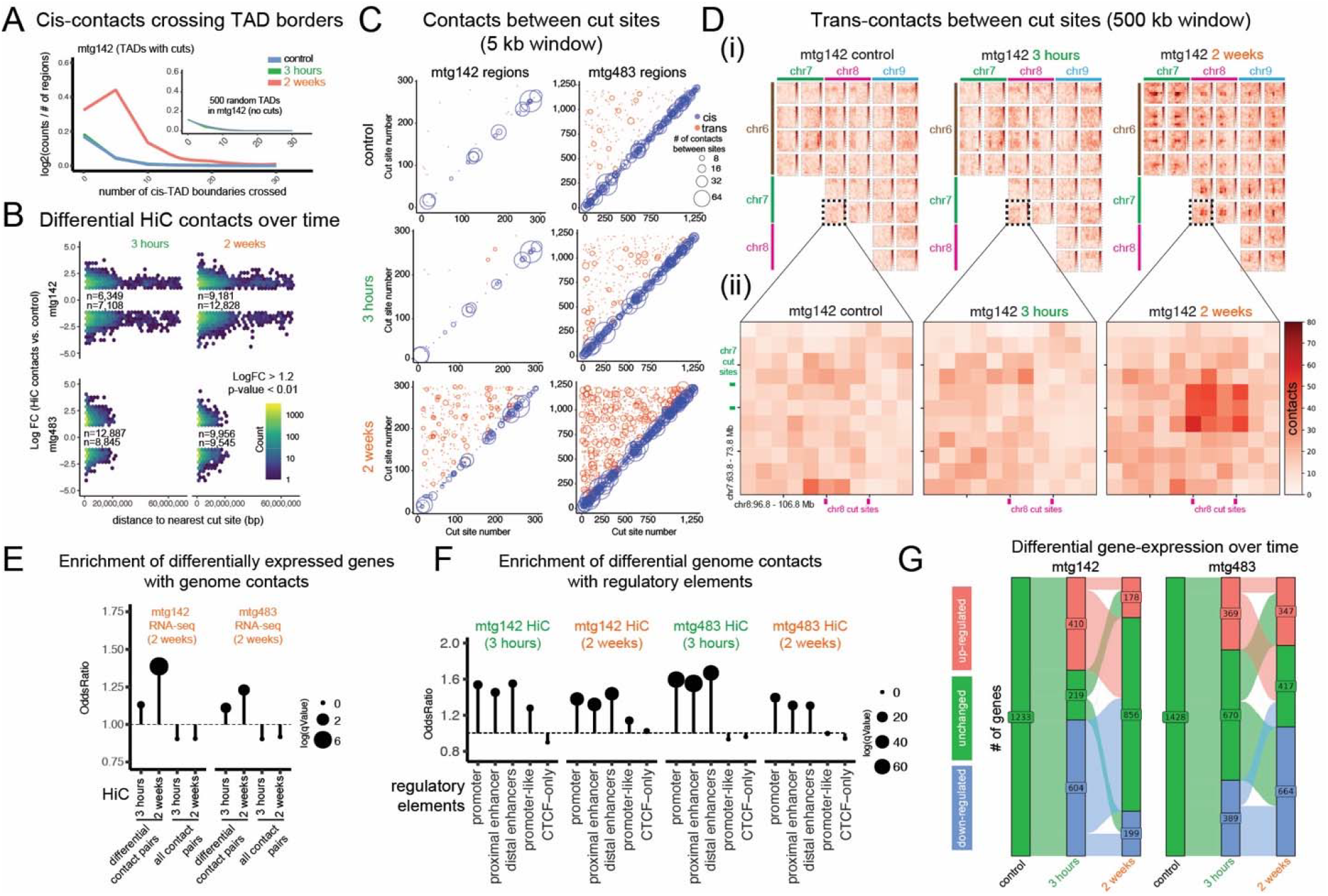
Reprogramming of long-range 3D genome contacts and regulatory elements following DNA damage over time are associated with persistent alterations in gene-expression. A. Quantification of cis-contacts from TADs with DSBs that cross TAD borders over time. Inset shows 500 random TADs that do not have DSBs for comparison. B. Differential cis-contacts ordered by distance to cut site show long-range (mega-base) alterations from the nearest cut site. *C. Cis* and *trans* contacts within 5kb of cut sites (cut-site to cut-site). The axes represent cut sites ordered by chromosome and position number (cut site versus cut site on either axes). D. Trans-contact heatmaps using 500kb trans-contact windows around cut sites. (i) Trans-contact heatmap-of-heatmaps (10 Mb regions) between cut sites for selected locations (mtg142 condition). (ii) Trans-contact heatmap between a single regions over time (10 Mb regions), which is highlighted with a dash-lined box in (i). E. Genomic enrichment analysis for differentially expressed genes with differential genomic contacts. G. Genomic enrichment analysis for differential genomic contacts with regulatory regions. F. Significant up and down-regulated genes over time. Alluvial plot shows the number of genes that are unchanged and remain differentially expressed as compared to control over time (LogFC >1.2, p-value <0.05).

Consistent with these findings, differential analysis of *cis* contacts showed significant alterations, including long-range contacts that persisted over time in cells expressing both guide RNAs **(Fig. 3B and fig. S7D)**. However, these analyses did not distinguish between new contacts that form between cut sites, alterations that could be important during the DNA damage response (*19*). Using a 5kb window and restricting our analyses to changes in contacts that occur only between any two cut sites, we demonstrate that chromatin interactions evolve over time, accumulating in an increase in both *cis* and *trans* contacts **(Fig. 3C and fig. S8A)**. In this analysis, *trans* contact changes were not as pronounced as *cis* contact changes, likely because these occur less frequently.

To visualize *trans* contacts more clearly, we expanded our window size to 500kb around cut sites and were able to identify the formation of significant *trans* contacts around damaged regions as shown in **Fig. 3D and fig. S9A to D**. These were less prevalent at 3 hours compared to the 2 week time point, emphasizing how chromatin organization evolves over time after the introduction of DSBs. Our data suggest that architectural disruptions can be actively induced by targeted DSBs alone, without an underlying genetic disorder. These findings echo recent discoveries of pathological chromatin rewiring in disease contexts such as fragile X syndrome and pediatric ependymoma. Specifically, BREACHes (Beacons of Repeat Expansion Anchored by Contacting Heterochromatin) (*20*) and TULIPs (ultra-long-range B compartment contacts) (*21*) were recently identified as large-scale 3D genome remodeling phenomena that emerge in disorders with intrinsic genome instability. These findings imply that DSBs may be sufficient to trigger a breach of chromatin domain boundaries and induce genome-wide reorganization.

To determine if changes in gene expression could be linked to changes in chromatin organization, we analyzed changes in interactions involving differentially expressed genes and found a significant enrichment at both 3 hours and 2 weeks in comparison to all contact pairs **(Fig. 3E)**. A similar analysis for regulatory elements revealed that promoters and enhancer sequences were enriched for differential genomic contacts **(Fig. 3F)**. Importantly, among the differentially regulated genes in the mtg142 and mtg483 conditions, 30.1% (n=377) and 70.8% (n=1,011) were persistently up or downregulated by the 2 week timepoint **(Fig. 3G)**. These findings indicate that the introduction of DSBs leads to transcriptional reprogramming in which a subset of gene expression changes persist until the 2 week timepoint. Transcriptional reprogramming is initially associated with a transient increase in intra-TAD contacts at 3 hours, but by 2 weeks there is a shift towards the formation of new longer-range, inter-TAD and *trans* contacts that correlate with increased A-B compartment mixing.

### Large structural variations accumulate over time and correlate with contact frequency and gene density and activity

DSB repair can lead to alterations in genetic sequence and less frequently, large structural variations (SVs), depending on the fidelity of the repair pathway involved. It is known that the emergence of SVs can be influenced by the non-genetic or chromatin context of the break (*19, 22, 23*), but this is typically challenging to study in a systematic fashion after the random introduction of radiation induced DSBs in individual cells. We capitalized on the uniformity of this Cas9-mediated DSB approach to examine the chromatin contexts of newly generated SVs after DNA damage induction.

We designed biotinylated DNA probes using complementary sequences that flanked mtg142 or mtg483 cut sites and performed high-coverage targeted sequencing to profile mutations **(fig. S10A)**. An inherent challenge in this process is that mtgRNAs by definition target the same sequence in the genome and there is some sequence degeneracy around the cut sites that limits the number of unique cross-compatible DNA probes that can be designed. Working within these constraints, we created two panels of 402 and 979 probes targeting 36% of mtg142 sites and 32% of mtg483 sites, respectively, with one to four probes flanking each cut site **(fig. S10B and data S5 to 6)**. As shown in **fig. S10C** we were able to detect small deletions and insertions at known cut sites, with sequencing depths with a median of ~1,100X after pooling replicates. As expected, nearly all cut sites had some level of detectible small indels, as highlighted in **fig. S10D**.

Large structural variations (SVs) were detected at nearly every sequenced cut site, with translocations, inversions, duplications and deletions seldom exceeding a frequency of 5% at any given cut site **(Fig. 4A)**. The total frequency of any large SV at a given DSB site is shown in **Fig. 4B and fig. S11A to B**, we did not detect any large insertions. Interestingly, we did not see any direct correlation between cut site efficiency (as assessed by isBLISS sequencing) and the frequency of SVs **(fig. S11C)**, suggesting that SV formation was influenced by other factors. Since these cut sites all shared very similar genetic sequences, we examined the local chromatin environment of the SV anchor sequences.

**Figure 4.**
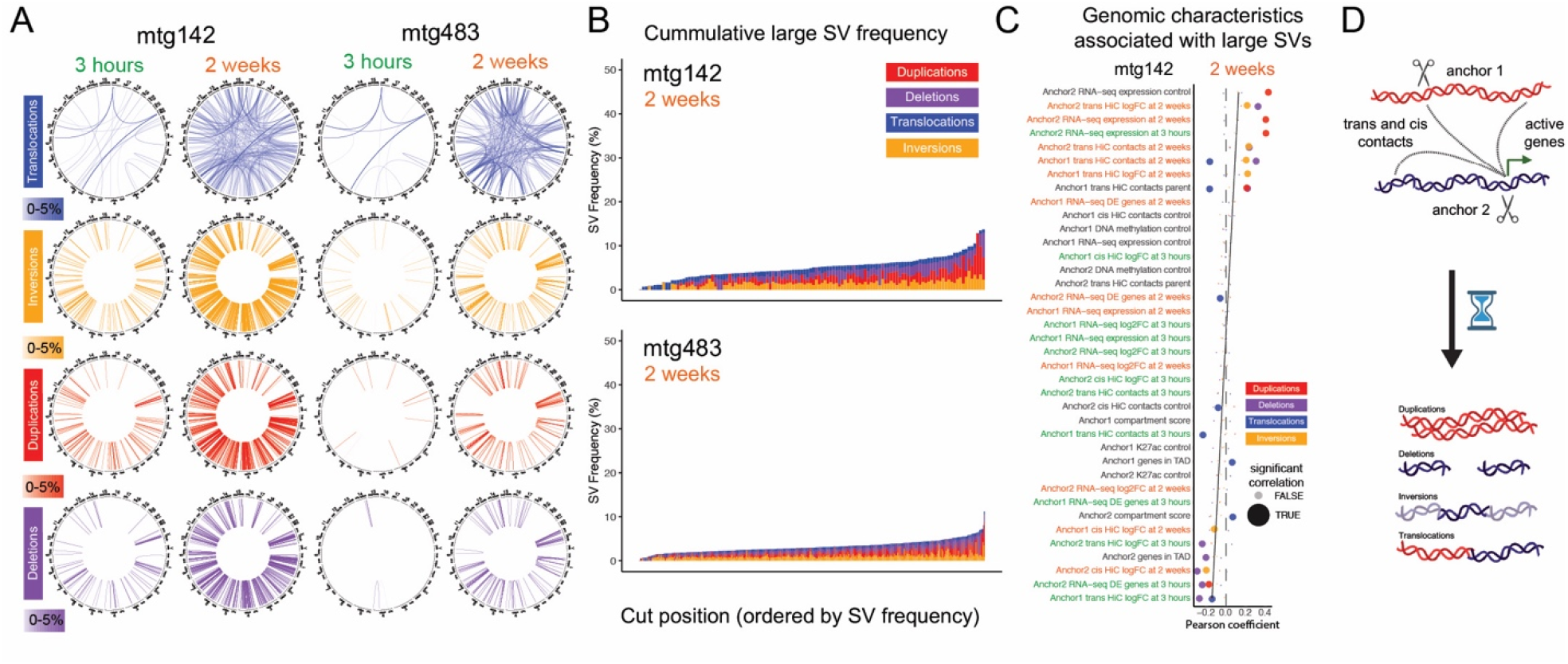
Large structural variations accumulate over time and correlate with contact frequency and gene density. A. Circular genomic plots representing large structural variations (SVs) over time between two cut sites (compared to control condition). Lines are shaded by locus SV frequency (0-5%) and colored by structural variation type. B. Cumulative large SV frequency (y-axis) ordered by total frequency along the x-axis. Each stacked bar represents a cut site. The two-week time points are shown. C. Correlation coefficients of genomic measurements performed at baseline (control), 3 hours and 2 weeks correlated to SVs that are detected at 2 weeks in condition mtg142. Anchor 1 and 2 demark the ends of the SV breakpoints (ordered numerically by chromosome number and location). Anchor specific genomic measurements (y-axis) are ordered by average Pearson correlation coefficient of the 4 different large SVs (x-axis). D. Scheme highlighting that large structural variations increase over time and correlate with contact frequency and gene density or activity following DNA damage.

For each SV category, we compared the genomic characteristics of surrounding regions of sequenced cut sites over time, and calculated correlation coefficients at either end of the SV (anchor 1 or 2) **(Fig. 4C and fig. S11D)**. This analysis included a complete cross correlation of the number of genes in the regions, RNA-seq, HiC, H3K27ac, DNA methylation and SV frequency **(fig. S12 to S15)**. Because alterations or shifts in or RNA-seq, HiC, H3K27ac and DNA methylation are distinct from high or low basal levels, we included both log fold-change and average levels at all timepoints as separate variables. Although our results show statistically significant correlations, no single genomic characteristic alone could strongly predict SV formation. However, a high frequency of *cis* or *trans* contacts and high gene-density or activity within one or both anchors emerge as characteristics associated with their emergence **(Fig. 4D)**.

## Discussion

In this study, we used pre-mapped DNA double-strand breaks (DSBs) in human glioblastoma stem cell lines to systematically track genetic and non-genetic changes over time. Our findings demonstrate that DNA damage can precipitate mega-base scale alterations in DNA and chromatin modifications, *cis* and *trans* genome interactions and change gene expression. Notably, these shifts remain detectable weeks after the initial DSBs, indicating that the cellular response to DNA damage can go far beyond the generation of transient repair phenotypes, with significant implications for cancer biology and normal cellular function.

While it is known that chromatin structure and gene expression change in the context of DNA damage stress (*2, 12, 24*), prior studies have focused on acute responses (*4, 13, 14*)— typically within hours of damage—or have examined long-term consequences in clinical samples without a means to associate observed chromatic and gene-expression changes to specific, traceable DSB events (*1, 2, 11*). Our data build on these studies, demonstrating the importance of evolving states that follow damage over time in cancer cells. Here we have shown that genomic reprogramming involves a transient gain of intra-TAD and interactions and the delayed formation of new long-range inter-TAD and trans contacts, that correlate with durable changes in DNA methylation, histone acetylation, gene expression and the emergence of large structural variations, all of which have important functional consequences.

More broadly, our findings suggest that the capacity of DSBs to induce 3D architectural reorganization may represent a unifying mechanism across both experimental and pathological contexts. While prior studies of fragile X syndrome and pediatric ependymoma have linked chromatin misfolding to disease phenotypes via structures like BREACHes (*20*) and TULIPs (*21*), our results indicate that reorganization can be context independent. This implies that chromatin architecture is not merely a passive reflection of sequence alterations or disease states but it can be actively reshaped by damage itself. The ability of targeted DSBs to provoke inter-compartment mixing and long-range contact remodeling suggests a generalizable vulnerability of genome architecture, with potential relevance for understanding how genome topology contributes to plasticity, repair, and cellular identity.

Although we examined widespread DSBs in a cancer and therapy context, DSBs occur during diverse normal cellular processes including gene transcription, development and signaling. For example, during lymphocyte development there are DSBs introduced during programmed recombination of antigen receptor loci, where the number of breaks introduced per cell is controlled to prevent widespread damage and genome instability (*25, 26*). Widespread DSBs may also occur during normal lymphocyte and neural development, where the number of DSBs depends on the specific context (*27, 28*). Post-mitotic neurons are also found to generate DNA damage with neuronal activation to enable the expression of early response genes critical for plasticity and memory formation, but it is not known whether these have persistent effects (*29*). Indeed, it may be possible that physiologic DSBs, or DNA damage in cancer cells, are similarly capable of inducing lasting alterations to promote genomic remodeling.

Despite these growing insights, several limitations and questions remain. First, consideration should be given to the type and variety of damage. Inducing DSBs via multi-target CRISPR–Cas9 represents a specific type of controlled, damage, whereas clinical DNA insults (e.g., from irradiation or chemotherapy), or random damage from genomic instability, are often more stochastic and can vary widely in, number, density, genomic context and type of insult. Such variability may accentuate or dampen genomic reprogramming, and future work comparing different damage models could clarify a threshold at which persistent changes become locked in. Additionally, while our focus centered on glioblastoma, it is unclear how broadly reprogramming events apply to other cancer types, disease states, or to nonmalignant cells, whose chromatin landscapes may respond differently to the introduction of widespread DNA damage.

Our data has broad implications for cancer biology as they suggest that cells sustaining mass DNA damage from genotoxic stress may adopt new chromatin configurations and gene expression programs that promote survival or drive therapy resistance. Understanding the precise pathways that stabilize reprogramming events could enable the development of therapeutic strategies aimed at preventing or reversing their occurrence to curb the emergence of resistant cell states. Overall, the findings presented in this study highlight DNA damage–induced chromatin rewiring as a potentially powerful mechanism underlying cancer cell plasticity and they underscore the need to integrate studies of DNA repair with a deeper exploration of the longitudinal effects of damage.

## Acknowledgments

FgH1tUTCherry was a gift from Marco Herold (received through Addgene plasmid # 85552). LentiCas9-Blast was a gift from Feng Zhang (received through Addgene plasmid # 52962). We thank the NYU Langone genomics core and NYU Langone microscopy for their services and technical assistance. We thank Integrated DNA Technologies (IDT) for their services and technical assistance.

## Author contributions

Conceptualization: ASM, JPC, JS, EPS

Methodology: ASM, CD, JPC, JS, EPS

Investigation: ASM, KC, CD, RE, ES, JK, YD, ZZ, MG, GC

Visualization: ASM, KC, CD

Funding acquisition: ASM, JS, EPS

Writing – original draft: ASM

Writing – review & editing: all authors

## Data and materials availability

All data, code, and materials used in the analysis will be available in some form to any researcher for purposes of reproducing or extending the analysis.

## Accession numbers

Pending

## Supplementary Materials

Materials and Methods

Supplementary Text

Figs. S1 to S14

Data S1 to S6

