## Supplemental File for "Persistent chromatin alterations and gene expression reprogramming follow widespread DNA damage in glioblastoma"

**The PDF file includes:**

Materials and Methods

Supplementary Text

Figs. S1 to S15

References

**Other Supplementary Materials for this manuscript include the following:**

Data S1 to S6

Materials and Methods

Cell culture and multi-target guide RNA expression

Patient derived glioma stem-like cultures (GSCs) were cultured in serum free media without doxycline to enrich for stem-like states, as detailed in a publication from (*1*). GSC20 was used for most mtgRNA experiments to compare timepoints and different muti-target guide RNAs. U87 glioma cells were utilized for clonogenic assays, they were grown in DMEM containing 10% FBS (doxycycline free). Lentivirus generation was performed using HEK 293T cells grown in DMEM containing 10% FBS.

Multi-target guide RNA sequences (including the PAM sequence) are as follows: mtg483: 5’-AGTGGGTGCAGCCCATGG, mtg142: 5’-GGCTTCCAGGTCACAGGT. Guide RNAs were cloned into 3^rd^ generation lentiviral vectors using standard techniques. The guide RNA was housed in a doxycycline inducible construct, FgH1tUTCherry (addgene #85552). To generate stable cell lines, they were initially infected with lentivirus containing human codon-optimized S. pyogenes Cas9 protein with blasticidin resistance (lentiCas9-Blast, addgene #52962) and selected with Blasticidin for 7 days. Following lentiviral transfection with mtg142-, mtg483- or empty-FgH1tUTCherry, cells were sorted via FACS and cultured further for downstream experiments. For all experiments, to induce expression of a mtgRNA, cells were treated with 10ng/uL of doxycycline for 3 hours, followed by two centrifugation washes with media to remove doxycycline at the 3-hour timepoint.

Immunofluorescence

Glioma stem cells were dissociated into single-cell suspensions and seeded onto coverslips pre-coated with laminin and poly-L-ornithine. Following the specified treatments, cells were fixed in 4% paraformaldehyde (PFA), then blocked with 2% bovine serum albumin (BSA). Immunostaining was performed with a primary antibody against γH2AX (Millipore 05-636), and cell nuclei were counterstained with DAPI. Coverslips were subsequently mounted and imaged by fluorescence microscopy.

Clonogenic assays

Human U87 glioma cells were seeded for clonogenic assays and either exposed to radiation or subjected to mtgRNA induction. Radiation was performed using a CellRad per manufacturer instructions. After one week of incubation, surviving colonies were fixed and stained with crystal violet. Each condition was performed in triplicate (n=3) to ensure reproducibility of the results.

In-suspension breaks labeling in situ and sequencing (sBLISS or isBLISS) and double-strand break definitions

Double-strand break sequencing in-suspension breaks labeling in situ and sequencing, or sBLISS or isBLISS) was performed as described in detail in Bouman et al. (*2*). Summarized here, the protocol begins with the crosslinking of cells with 2% PFA. Cells are lysed to expose the DSB ends, which are subsequently blunted *in situ* and ligated to the sBLISS adaptor. Genomic DNA is then extracted and sonicated. Library preparation proceeds with *in vitro* transcription for linear amplification of the template. The library is then ligated to a new adaptor and annealed to a complimentary oligonucleotide to enable reverse transcription of each template. The entire library is amplified with sequencing primers that include indexes and unique barcodes. Following quantification and quality control checks via tapestation analysis, the libraries are sequenced via an Illumina NovoSeq6000 with 10% PhiX spiked in.

Raw sequencing data was first processed by author instructions. Briefly, the blissNP pipeline (*3*) was used to generate a BED file that averaged both replicates with counts of every unique double-strand break that had been sequenced. Signals from the isBLISS data were normalized to achieve a mean signal of 1 across all bases. We calculated the total signal by multiplying each region’s length by its signal value and divided by the total number of bases to determine the current average signal. We then applied a normalization factor to each region’s signal, bringing the average signal across each sample per base to 1.

To define the final set of mtgRNA-CRISPR-cas9 cut sites we defined a potential ‘universe’ of theoretical guide-RNA landing sides that includes a perfect match (0 mis-matches to the genome with the guide RNA sequence), 1 mis-match (1 nucleotide is not matched) or 2 mis-matches (2 nucleotides do not match), up to two RNA bulges on the genome and up to two DNA bulges on the guideRNA. This universe of potential target sites was intersected with the normalized isBLISS signals to only include sites that had a double-strand break, yielding 339 cut sites for mtg142 and 1,386 sites for mtg483. Cut sites were visualized either directly by IGV (version 2.18) or by circlize (version 0.4.16) in R (version 4.1.2).

Transcriptomic profiling

RNA was extracted using the AllPrep genomic DNA/RNA extraction kit (Qiagen). Experiments were performed in duplicates (n=2). RNA was subject to tapestation for RIN quality assessment with a minimum quality score of 9 required to proceed with library preparation. RNA libraries were prepared with TruSeq Stranded Total RNA Gold for whole-transcriptome sequencing according to manufacturer protocols. In brief, unwanted rRNA was depleted, mRNA/cRNA was converted to double-stranded cDNA, end-repaired, adapter-ligated, and PCR-amplified. The final libraries were purified, quantified, and assessed for integrity prior to sequencing with a tapestation. Sequencing multiplexed and performed on a NovoSeq 6000 with a goal of achieving at least 75 million reads per a sample.

For data processing, adapters along with low-quality bases were trimmed using Trimmomatic. The resulting reads were aligned to the human reference genome (hg38) with STAR, and additional alignments were performed against other species and common contaminants for quality control. Normalized genome browser tracks were generated to visualize coverage, and the 5′/3′ bias was assessed with Picard. Gene-level counts were then summarized to create a genes-by-samples matrix, and differential gene expression analysis was conducted using standard DESeq2 (version 1.46.0) procedures.

DNA-methylation profiling

DNA was extracted using the Qiagen AllPrep genomic DNA/RNA extraction kit (Qiagen). Experiments were performed in duplicates (n=2). Genomic DNA was processed using the TruSeq Methyl Capture EPIC Library Prep Kit (Illumina) according to the manufacturer’s protocol. Briefly, high-quality DNA (as determined by tapestation analysis) was fragmented via a Covaris S220, followed by end-repair, 3′ adenylation, and ligation of methylation-compatible adapters. The resulting fragments underwent bisulfite conversion to convert unmethylated cytosines to uracils, while preserving methylated cytosines. The libraries were then hybridized to the TruSeq Methyl Capture EPIC probes for targeted enrichment of methylated regions, followed by bead-based capture. Finally, the enriched libraries were PCR-amplified, purified, and assessed for quality with a tapestation prior to sequencing. Sequencing multiplexed and performed on an Illumina NovoSeq 6000.

Partial genome bisulfite sequencing data from TruSeq Methyl Capture was trimmed for adapters and low-quality bases using Trimmomatic, then aligned to the bisulfite-converted hg38 reference genome via Bismark and Bowtie2. Duplicate reads were removed in Bismark, and methylation calls were subsequently extracted. Further processing was conducted in methylKit, beginning with coverage-based filtering to remove bases below 10× coverage and above the 99.9th percentile. Samples were merged and used to calculate percent methylation. Differential methylation analyses were performed on the merged data, applying thresholds for q-values (e.g., <0.01) and methylation differences (≥25%), and the resulting differentially methylated bases or regions were stored for further downstream interpretation.

Chromatin Immunoprecipitation

ChIP–seq was performed essentially as described in (*4*) using a dual crosslinking method and *in situ* tagmentation. In brief, ~1 million cells per replicate were first crosslinked in 1.5 mM ethylene glycol bis(succinimidyl succinate) (EGS) for 30 min, followed by 1% formaldehyde for 10 min, then quenched with 125 mM glycine. After washing, cell pellets were snap-frozen in liquid nitrogen and stored at −80 °C. Pellets were lysed in the presence of protease and deacetylase inhibitors, and the chromatin was sonicated to the desired size range. Drosophila spike-in chromatin (Active Motif) was added at 2% by mass for normalization, and immunoprecipitation was carried out with protein A magnetic beads (ThermoScientific #10001D) conjugated to H3K27ac (2.5uL per 1x10^6^ cells) (Abcam #ab4729), or spike-in antibody (2% of ug of target Ab, Active Motif #104597). After extensive washing, *in situ* tagmentation was performed on bead-bound chromatin to incorporate sequencing adapters, and crosslinks were reversed by overnight proteinase K treatment. DNA was purified by column-based methods, then PCR-amplified to produce the final libraries. Libraries were size-selected with AMPure beads, checked for concentration and fragment distribution (Qubit and Tapestation), and quantified by qPCR prior to multiplexed sequencing on an Illumina NovaSeq 6000.

Genomic enrichment analysis

To perform enrichment testing, we performed genomic enrichment analysis using a Locus OverLap Analysis (LOLA) (*5*). This method uses a Fisher’s exact test with false discovery rate correction to see if segments of the genome marked by differential HiC contacts, or gene-expression, are statistically enriched over other parts of the genome in a pairwise manner. Cis-regulatory element defintions are defined by ENCODE (*6*). Processing of all other genomic measures (RNA-seq and HiC contacts) are described elsewhere in the methods section.

Genomic contact (genome-wide chromosome conformation capture) sequencing and data processing

For HiC (genome-wide chromatin contact conformation), the ARIMA Standard Hi-C Kit was used following manufacturer instructions. In brief, cells were crosslinked with 2% formaldehyde for 10 minutes at room temperature, and the reaction was quenched with glycine. Chromatin was digested with the restriction enzyme mixture provided in the ARIMA Standard Hi-C Kit, followed by a biotin fill-in step and proximity ligation according to the manufacturer’s protocol. Experiments were performed in duplicates (n=2). After reversing crosslinks and purifying the DNA, adapters were ligated to the fragment ends, and library amplification was performed. Sequencing multiplexed and performed on an Illumina NovoSeq 6000.

Standard pipelines were used to process cis-genome HiC contact data unless noted otherwise. HiC Bench was used in conjunction with standard bioinformatics tools to ensure reproducibility. Alignment was performed with bwa (version 0.7.17) using settings -A1 -B4 -E50 -L0. After alignment, reads were filtered with samtools (version 1.3) and gtools (version 3.0) using a mapping quality threshold of 20, no minimum distance, and a maximum offset of 500. Duplicate reads were also removed during this step. Filtered contact pairs were converted to hic format using juicer (version 1.5) and normalized using Knight-Ruiz (KR) normalization. These were then converted to cool format using hicexplorer (version 3.7.2) in various resolutions for further visualization and processing in GENOVA (version 1.0.0).

HiC data visualization with GENOVA using cool file outputs included the following steps. Centromeres were masked from analysis. Insulation scores for Topologically Associated Domain (TAD) inference was called using hic resolutions of 20kb with replicates combined. Compartments were called using hic resolutions of 500kb using combined replicates and signing of compartments was determined using H3K27ac ChIP-seq data. Interaction scores between arms of chromosomes were visualized in saddle plots and data was exported for quantification.

Data processing of *trans* chromatin interactions from HiC data

*Trans*-chromatin contact maps were constructed at 1 Mb resolution separately for each replicate using Hi-C valid pairs from HiC-Pro (version 3.0.0) output. HiC-Pro processing options were as follows: ligation/digestion sequences (GATCGATC, GANTGATC, GANTANTC, GATCANTC), insert size range 100 to 800 bp, min and max fragment sizes set to 10 and 1,000,000, chromosomes XY were excluded and hg38 was used as the reference genome. Time point replicates were merged into a single chromatin map to reduce sparsity and enhance detection of *trans* interaction frequencies. Regions containing outliers were removed, and the contact maps were balanced using the Knight-Ruiz algorithm with default parameters (*7*). Subsequently, the contact maps were scaled using Hi-C contact distance-dependent median-of-ratios size factors to bring all samples to the same scale and enable direct comparisons (*8*). For each pair of cut sites, *trans* contact maps were visualized with a 5 Mb window on either side. For downstream analysis, the matrix corresponding to the *trans* contact map with a 1 Mb window on either side was extracted. Summation and averaging of the matrices were performed, followed by transformation into a one-dimensional vector. Finally, p-values were calculated using the Mann-Whitney U test.

DNA probe design and pull down for deep sequencing of mtgRNA target sites

Two custom biotin-tagged capture panels (xGen DNA panels, Integrated DNA Technologies) were designed to enrich specific DNA regions, each featuring 120 bp probes with one to four probes flanking each targeted cut site, see **Supplemental Table S5-6**. One panel contained 402 probes (mtg142) and another contained 979 probes (mtg483). The degeneracy of genomic sequences around cut sites prevented the design of probes with high enough specificity to target all sites, therefore only probe/site combinations which could be targeted without significant overlap were included in the panel. Following library construction, the prepared libraries were hybridized with the panels according to the manufacturer’s guidelines, subjected to stringent washes to remove nonspecific fragments, and subsequently PCR-amplified. The enriched libraries were purified and assessed for quality with a tapestation prior to sequencing. Sequencing was multiplexed and performed on a MiSeq 2000 with 10% PhiX spike-in.

Small mutation and large structural variant calling:

To assess the frequency and distribution of small insertions and deletions (indels) and large structural variants (SVs) arising from CRISPR–Cas9-induced DNA double-strand breaks, we used separate, tailored analysis pipelines based on the sequencing assay. In both analysis pipelines, we constricted the analysis to verified cut sites and the pulled down regions that achieved coverage levels exceeding 100X. ENCODE defined blacklisted regions were filtered out. For small indel analysis, we applied the crispr-DART (Downstream Analysis and Reporting Tool) pipeline (9). Paired-end Illumina reads were subjected to quality control using FastQC (version 0.11.7) and trimmed with Trim Galore, followed by alignment to the human reference genome (hg38) using BBMap. To enhance detection sensitivity, reads overlapping putative indels were realigned using the Genome Analysis Toolkit GATK (version 3.8.0) for local realignment around indels and base quality score recalibration. Indel frequency profiles were visualized using IGV (version 2.18) and aggregated into heatmaps with deeptools (v3.5.1) using BigWig files output by crispr-DART.

For large SV analysis, we followed a parallel preprocessing workflow. Raw paired-end reads underwent quality control using FastQC (version 0.11.7) and adapter trimming via Trimmomatic (version 0.36). Reads were aligned to hg38 with BWA-MEM (version 0.7.17), and duplicates were removed using sambamba (version 0.6.8). BAM files were processed with GATK (v3.8.0) for local realignment around indels and base quality score recalibration. These realigned and recalibrated BAMs were used for structural variant calling with DELLY (version 0.8.1). DELLY identified deletions, duplications, inversions, and translocations using split-read and paired-end evidence from tumor-versus-control BAM comparisons. Output .bcf files were converted to .vcf using bcftools, and high-confidence SVs were annotated and visualized using Circos and other genome plotting tools.

Correlation of structural variant (SV) relationships with genomic features

Pearson correlation coefficients were calculated between well covered cut-sites from DNA-pull down sequencing. Large Structural Variation (SV) frequencies and 58 genomic features across TADs spanning timepoints with cis-chromosomal contacts (Hi-C), RNA-seq attributes, histone modifications (H3K27ac), and DNA methylation attributes (processed as described in a separate section). For each *trans*-contact genomic region with valid data (a sequenced cut-site), we calculated the mean Log2 fold-change across all overlapping observations at each time point. One-sample t-tests (comparing the sample mean to zero) were then performed using the stats (version 3.6.2) package in R, yielding a p-value for each region/timepoint combination. We applied the Benjamini–Hochberg (BH) correction to control the false discovery rate (FDR).

Data were analyzed separately for two independent conditions (mtg142 and mtg483) at three timepoints (baseline, 3 hours and 2 weeks post-treatment). For each SV type (deletions, duplications, inversions, insertions, and translocations), correlations were computed using pairwise complete observations to handle missing values. Statistical analysis was performed using R (version 4.1.2). Statistical significance was assessed using correlation tests with p-values adjusted for multiple comparisons (significance threshold p < 0.05), and insignificant correlations were masked in the visualization as noted in the figures. To identify patterns of relationships between genomic features and structural variant frequencies, we employed hierarchical clustering with complete linkage to organize correlation matrices. The resulting correlation matrices were clustered using a distance metric defined as d = sqrt(2(1-r)), where r is the correlation coefficient, producing a dendrogram-based organization of features. This clustering approach grouped features with similar correlation patterns together.

Correlation of genomic features between genomic regions

To compare and correlate changes in multiple types of genomic data (RNA-seq, ChIP-seq, DNA methylation, HiC genomic contacts), the genome was first segmented into regions defined by Topologically Associated Domains (as defined elsewhere in this methods section). Genomic features were averaged within that domain and average-log-fold change was calculated per region with statistics performed to assess significance, this was done as follows:

RNA-seq differential expression results (genes with log2 fold changes and p-values) were reduced to their midpoints to facilitate TAD overlap identification. For each TAD, all genes fully or partially residing within it were identified, and a mean log2 fold change was calculated across those genes per region. We used Fisher’s method to combine individual gene-level p-values into a single p-value for each TAD. Briefly, all gene-level p-values belonging to the TAD in question are converted to their natural logarithms and summed. The test statistic is then defined as −2 times this summed log(p). Under the null hypothesis, this statistic follows a chi-square (χ2) distribution with degrees of freedom equal to 2 × (number of p-values). The final combined p-value is calculated as the upper-tail probability of observing a chi-square statistic at least as large as the one computed.

H3K27ac (ChIP-seq) normalized bigwig files were converted to bedgraph format and an averaged region score was extracted per TAD region. Similarly, DNA methylation was averaged across TAD regions. Per-condition comparisons for ChIP-seq and DNA methylation data were then tested using a linear model framework in the limma package (version 3.62.2) using the fitting the model lmFit; relevant contrasts (e.g., “treatment vs. vector”) were defined for each experimental group, and empirical Bayes shrinkage was applied (eBayes). This approach yielded estimates of log fold-changes and corresponding p-values for regional comparisons.

To identify differential changes in cis-contact frequency by TAD regions, a DESeq2 (version 1.46.0) analysis was performed, fitting a negative binomial model with timepoint as the main factor. Empirical analysis of variance was done with DESeq, and results were extracted for each contrast of interest (3 hours vs. control, 2 weeks vs. untreated). Log2 fold changes, p-values, and adjusted p-values (via Benjamini–Hochberg correction) were extracted by genomic coordinates.

Supplementary Text

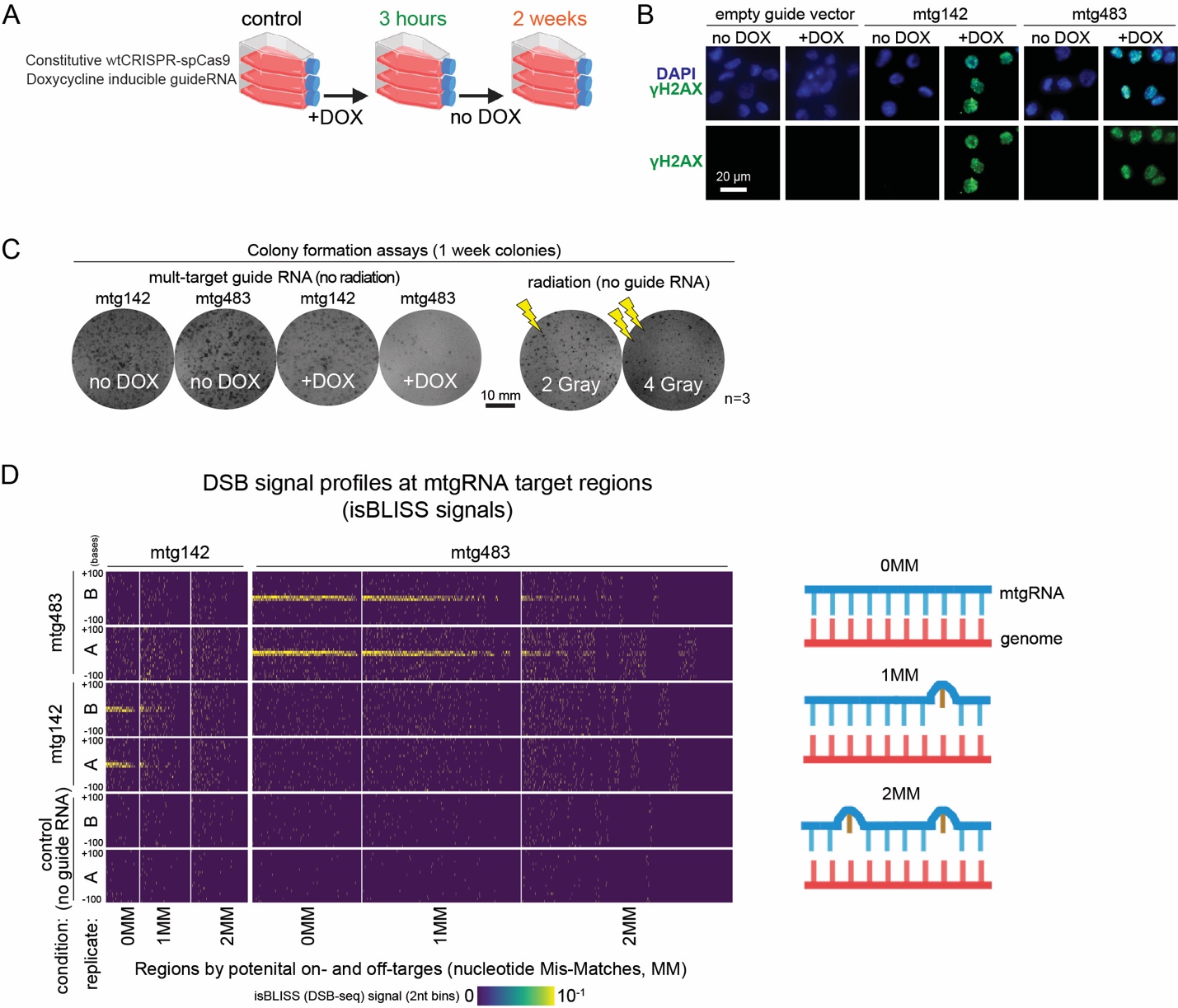

Fig. S1. Experimental design and quality control for the multi-target guide RNA system in Figure 1.

A) Human (patient derived) Glioma Stem Cells (GSCs) are stably transfected with constitutive wt-spCas9 and a separate doxycycline-inducible guideRNA. Control cells are treated with doxycycline for three hours for the first time point. After doxycycline withdrawal, the cells are allowed to recover, and a 2 week time point is collected.
B) Immunofluorescence staining of γH2AX three hours after doxycycline addition (+DOX) or no doxycycline addition (no DOX).
C) Colony Formation assays in U87 glioma cells comparing overall colony formation ability one week after induction of mtg142 or mtg483, or irradiation.
D) Heatmap of isBLISS signals (double-strand break sequencing). Rows represent sample condition (control, mtg142 and mtg483) and replicates (A or B). Columns represent regions in the genome that perfectly match (0MM) the guide RNA, or have 1-2 mismatches (1MM and 2MM).

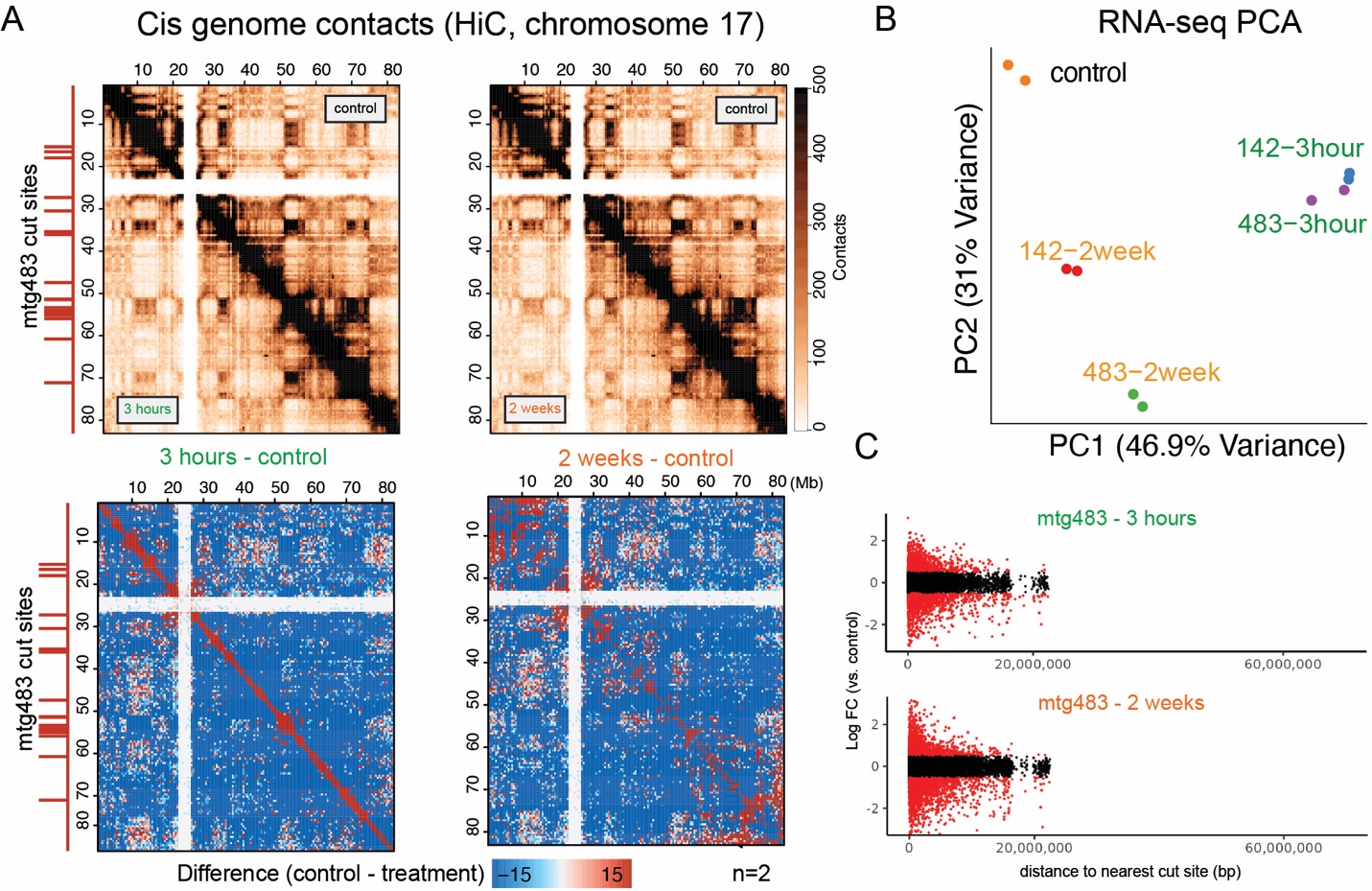
Fig. S2. Additional genome cis-contact maps and supplementary RNA-seq data for Figure 1.

A) Hi-C (genome-contact) maps of chromosome 17 showing contacts (top panel) and contact changes (bottom panel) over time (3 hours and 2 weeks). Cut sites for mtg483 for this region are shown on the left y-axis in red.
B) PCA of RNA-seq by replicate.
C) RNA-seq of differentially expressed genes marked as a function of distance to the nearest cut site for mtg483.

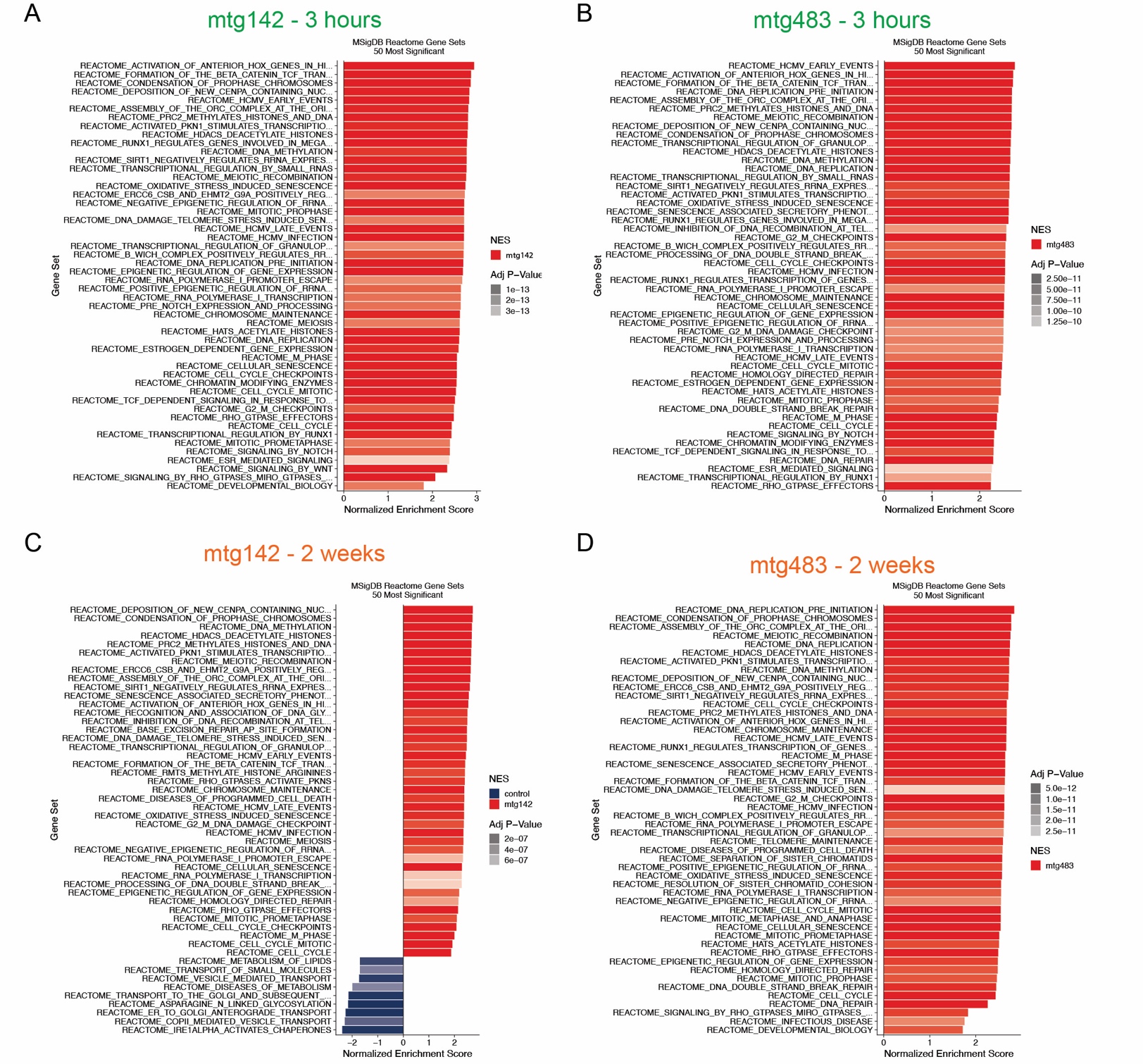
Fig. S3. Gene-Set Enrichment Data for Figure 1.

A) – D) Gene Set Enrichment analysis using the MSigDB REACTOME gene sets. A complete list of pathways, scores and genes are available in the Supplementary Data Tables.

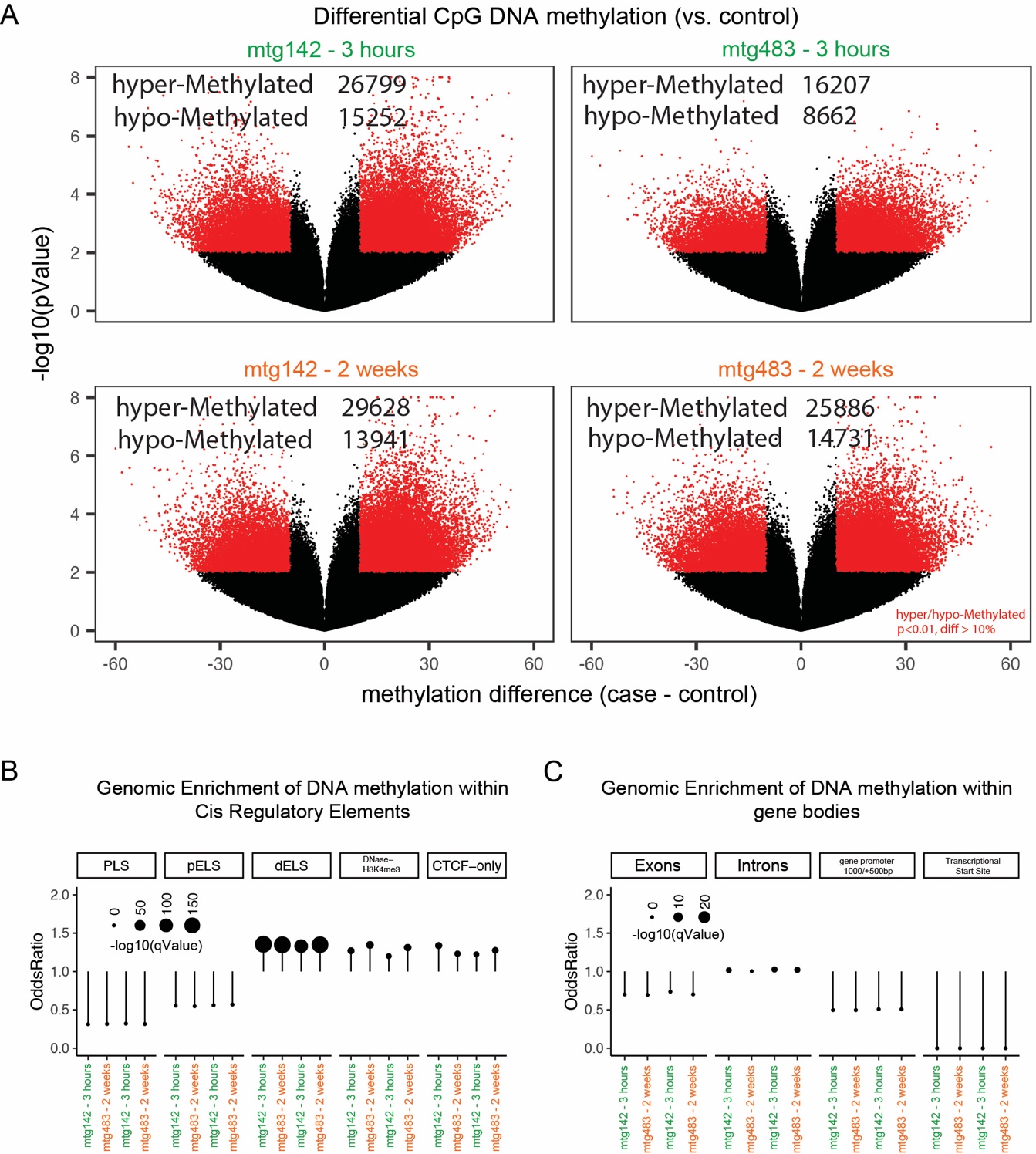

Fig. S4. Supplementary DNA methylation differential analysis and genomic enrichment

A) Differential DNA (CpG motif) methylation (sample vs. control condition).
B) Genomic Enrichment q-values for differentially methylated DNA at cis regulatory elements in the genome. The following abbreviations were used for ENCODE candidate Cis-Regulatory Elements (cCREs): Promoter-Like Sequence (PLS), proximal Enhancer-Like Sequence (pELS), distal Enhancer-Like Sequence (dELS).
C) Genomic Enrichment q-values for differentially methylated DNA at gene body locations across the genome.

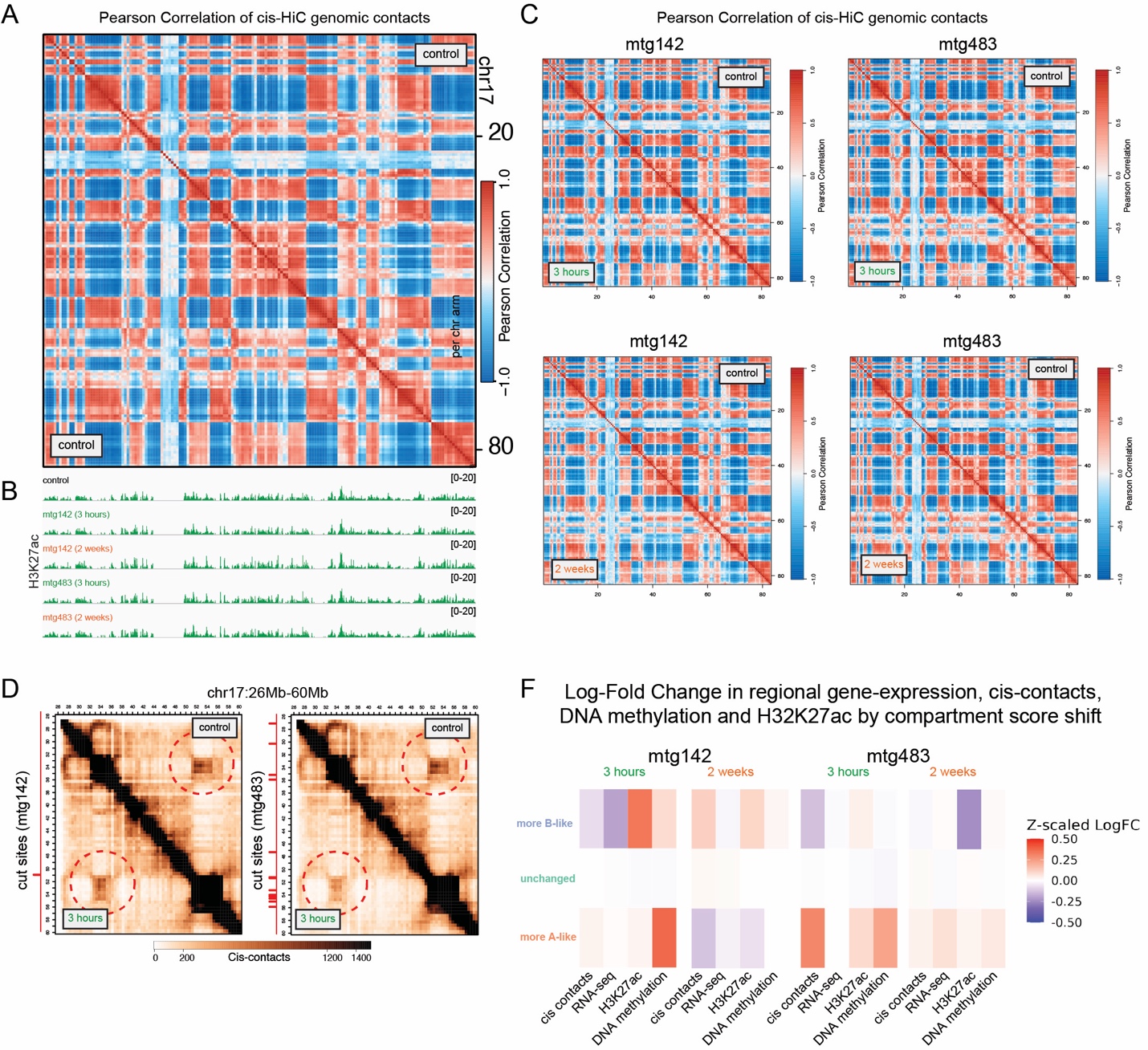

Fig. S5. Supplementary Data for Figure 2 showing Pearson Correlation plots and data used for compartment scoring and genomic changes averaged over TAD regions by degree of compartment shift.

A) Pearson Correlation of cis-HiC genomic contacts used to define compartments. Control is shown on both sides of this plot.
B) H3K27ac ChIP-seq signals across chr17, aligned with the Pearson Correlation plot in (A). These signals were used to define the direction of compartment scores.
C) Per-sample HiC Pearson Correlation plots of chr17 of mtg142 and mtg483 over timepoints (control top right of the diagonal, condition bottom left).
D) Genome contact maps (HiC) (control vs. 3 hours). Red circles highlight significant changes in long-range cis contacts near areas of cut sites.
E) Heatmap of z-scaled log-fold change of the averaged regional log-fold change in cis-contacts, gene-expression, DNA methylation and H3K27ac (columns), organized by degree of compartment score shift (rows). More A- or B-like shifts were defined by a change in negative or positive changes in compartment score >0.2). Regions were defined by TADs.

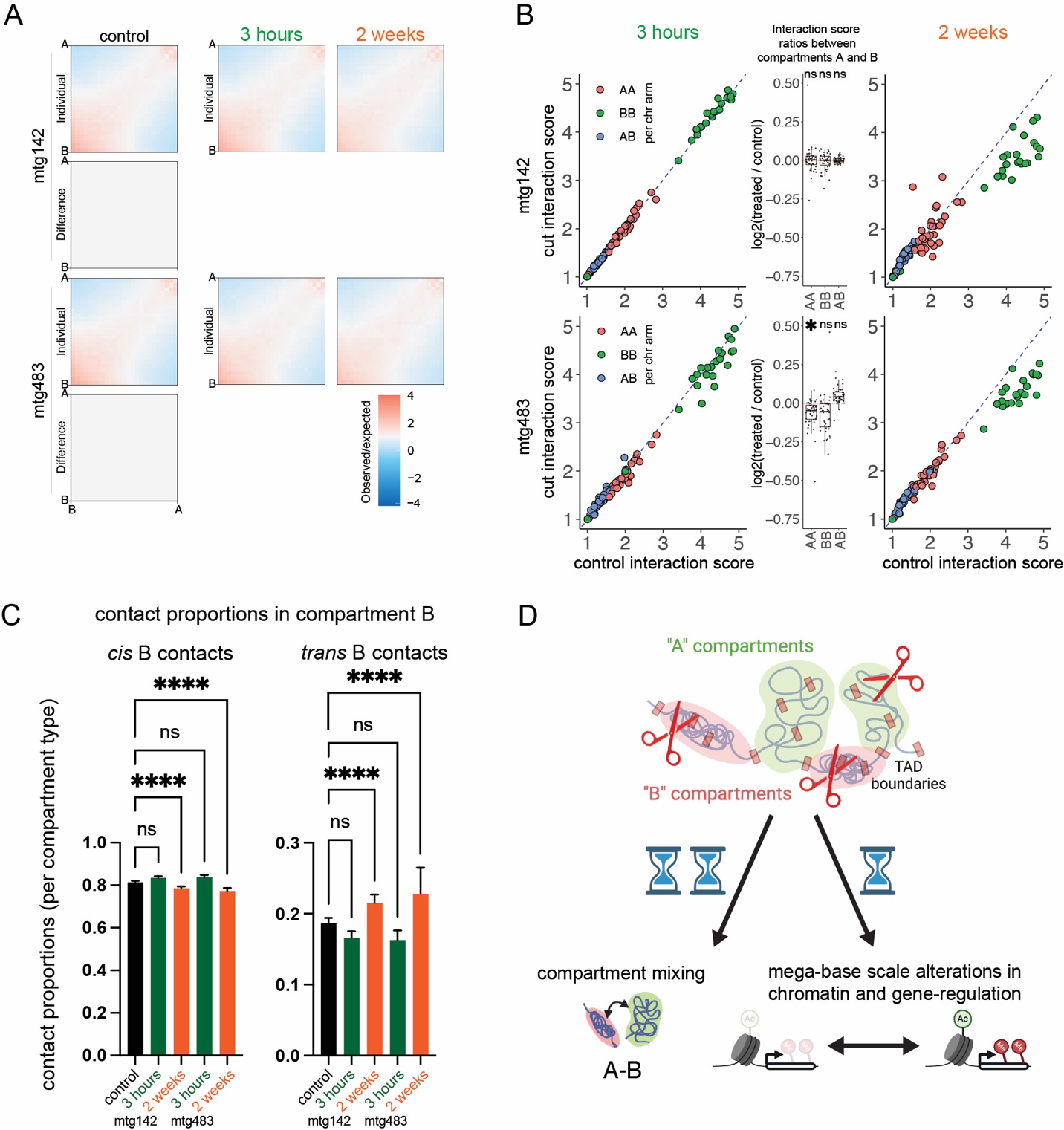

Fig. S6. Supplementary Data for Figure 2 showing interactions between compartment types and proportions of *cis* and *trans* contacts in B compartments.

A) Interaction score plots between all A and B compartments by condition and timepoint. Difference plots for mtgRNA minus control are in the main figure.
B) Scatter plots of interaction scores (by chromosome arm) with boxplots representing quotients (treated divided by control) to the right of each scatter plot (two-tailed one-sample t-test, * p-value < 0.01, ns = not significant). 2 week time point boxplots are in the main figure.
C) Contact proportions binned into *cis* or *trans* found in A compartments (ANOVA and Dunnett’s test, **** pValue < 0.0001, ns = not significant).
D) Schematic representation of increased A/B contacts, alterations in H3K27ac marks and DNA methylation over time.

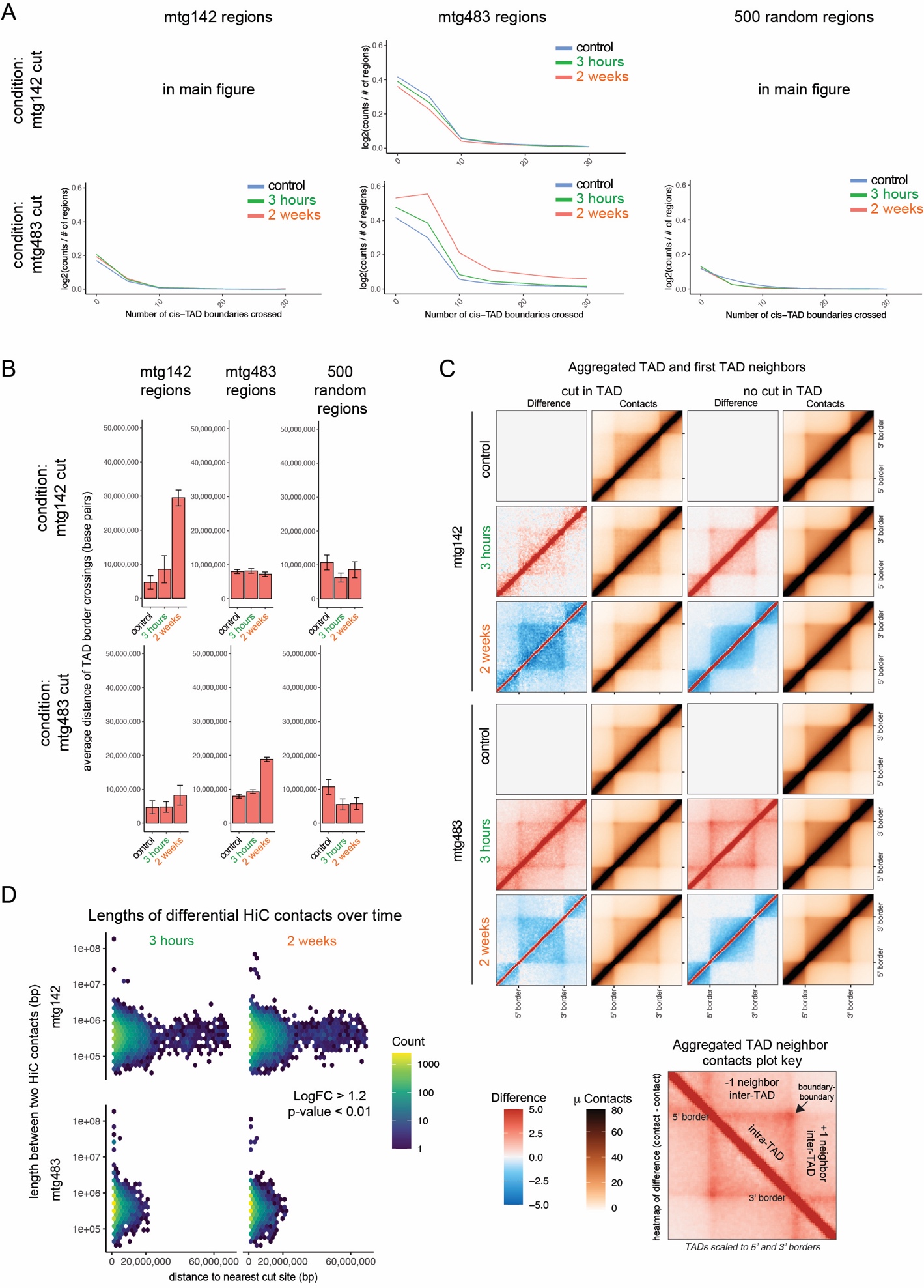

Fig. S7. Supplementary Data for Figure 3A-B showing cis-contact TAD crossings, aggregated TAD contact maps and lengths of differential contacts

A) Quantification of cis-contacts from TADs with DSBs that cross TAD borders over time. Columns (regions), represent the TADs what overlap with the indicated regions. Rows (condition containing mtg142 or mtg483) represent which mtgRNA was used. As a control, 500 random TADs that do not have DSBs were included, and respective regions were included (i.e. the mtg483 regions are shown in the second column for the condition that had mtg142 expressed).
B) Cis-contact distances of contact pairs that cross TAD borders (quantification of contacts from **(A)**.
C) Aggregated cis-contact maps and contact differences scaled to the 5’ and 3’ end of TADs. Each region represents a single TAD flanked by an upstream and downstream TAD. Scales and plot key are located at the bottom of the plots.
D) Contact lengths (y-axis) of differentially altered cis-contacts demonstrating the quantity and distribution of long-range contacts between conditions over time by distance from the nearest cut site (x-axis) (LogFC > 1.2 and p-value <0.02).

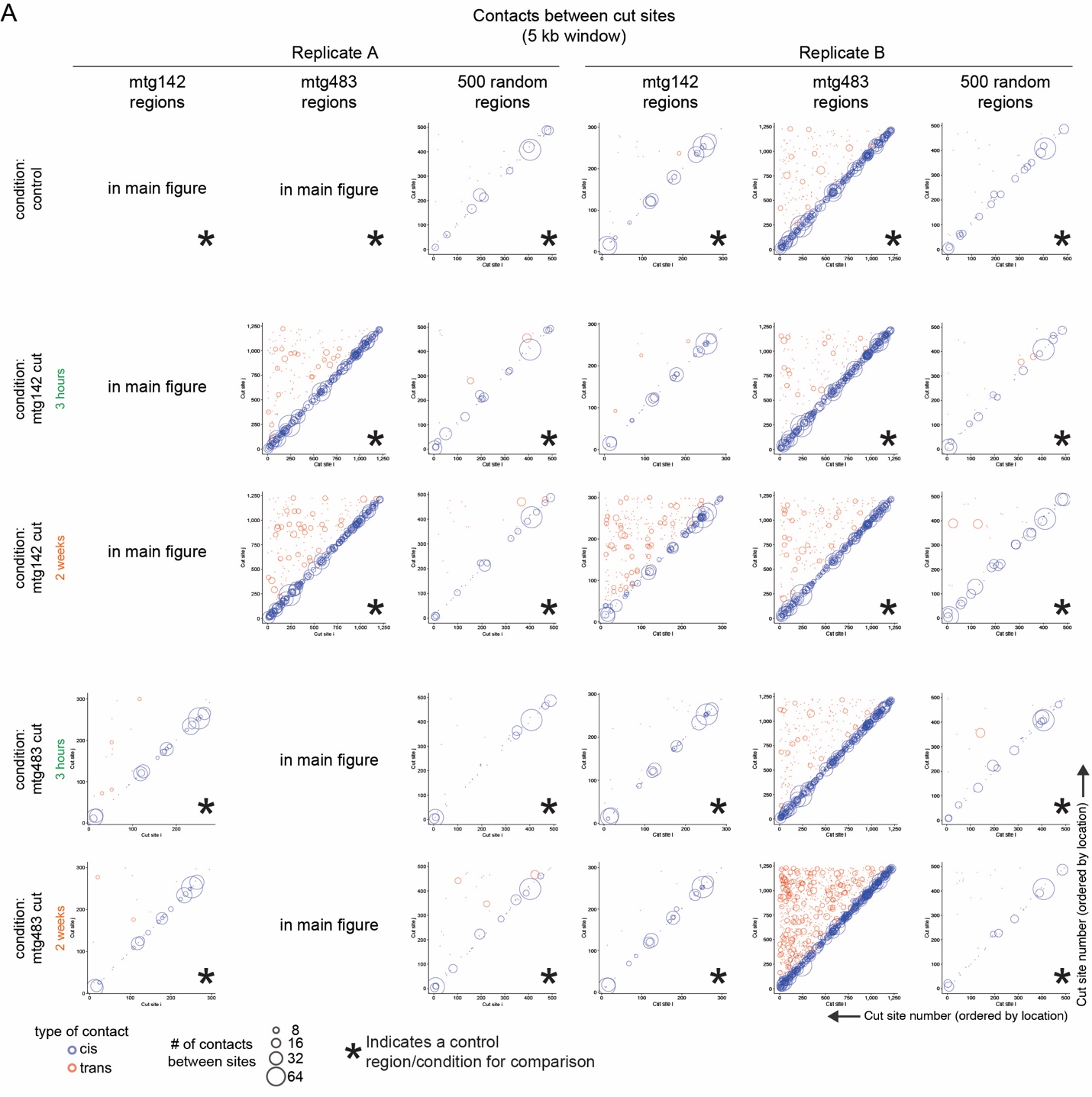

Fig. S8. Supplementary Data for Figure 3C

A) *Cis*- and *trans*-contacts within 5kb of cut sites (cut-site to cut-site) by replicate. The axes represent cut sites ordered by chromosome number (cut site vs. cut site). Data is organized by columns which represent regions used in the analysis, while rows present the experimental condition and timepoint. *Cis* and *trans* contacts are demarked by color, the size of each circle represents the number of contacts between two cut sites. An asterisk represents region/condition combinations shown as an experimental/analysis control. As a control, 500 random TADs that do not have DSBs were included, and respective regions were included (i.e. the mtg483 regions are shown in the second column for the condition that had mtg142 expressed).

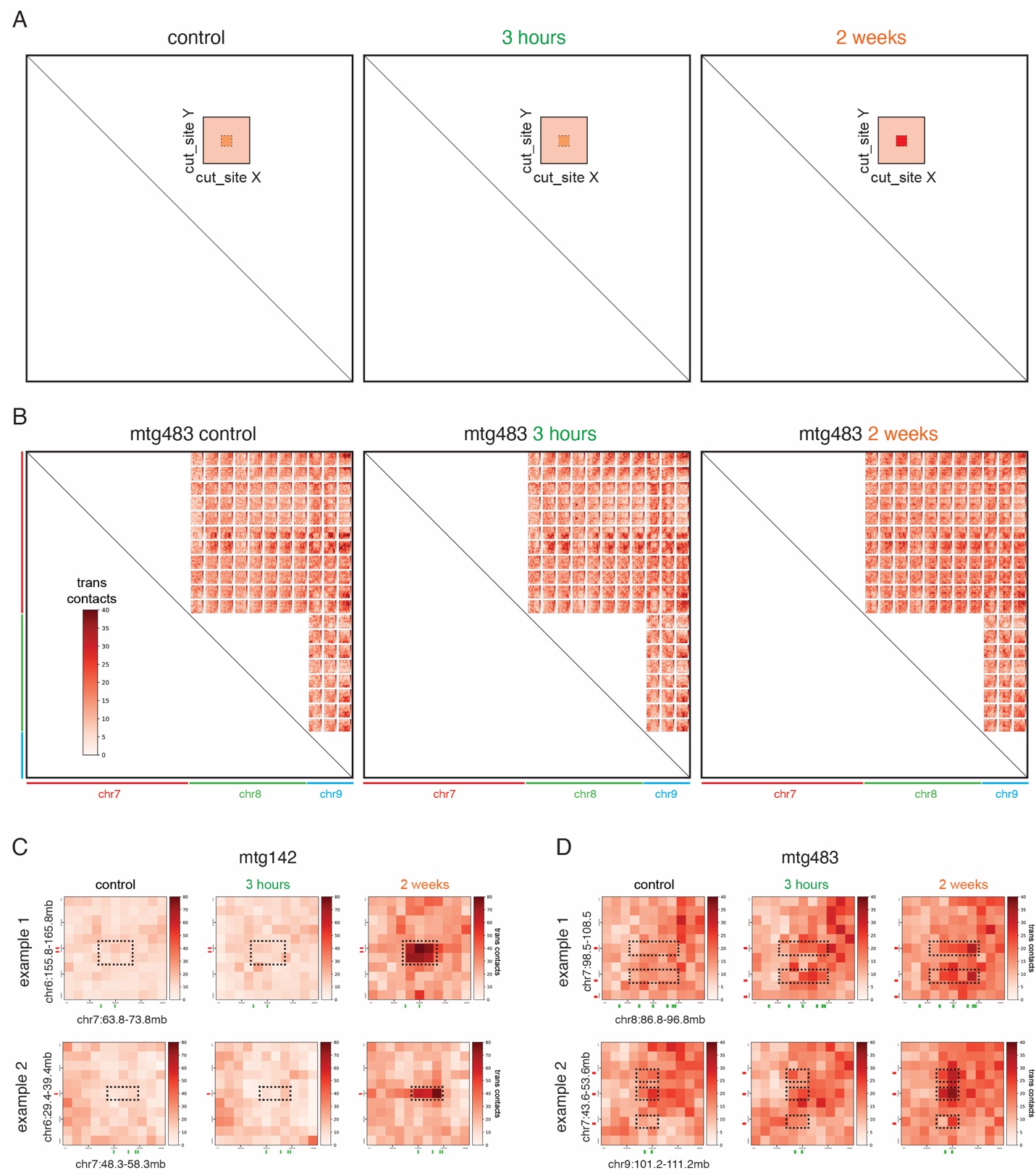

Fig. S9. Supplementary Data for Figure 3D – *trans* contact heatmaps between cut sites using 500kb *trans*-contact windows

A) Key diagram depicting that heatmaps are between two cut sites (x and y-axis) and that they are centered on a cut site (for Figure 3D and Supplementary Figure S8B-D).
B) *Trans*-contact heatmaps using 500kb *trans*-contact windows around cut sites (10Mb regions) between cut sites for selected locations (mtg483 condition).
C) and **D)** Representative *trans*-contact heatmap between single regions over time (10Mb regions). Selected areas with alterations are highlighted with a dash-lined box.

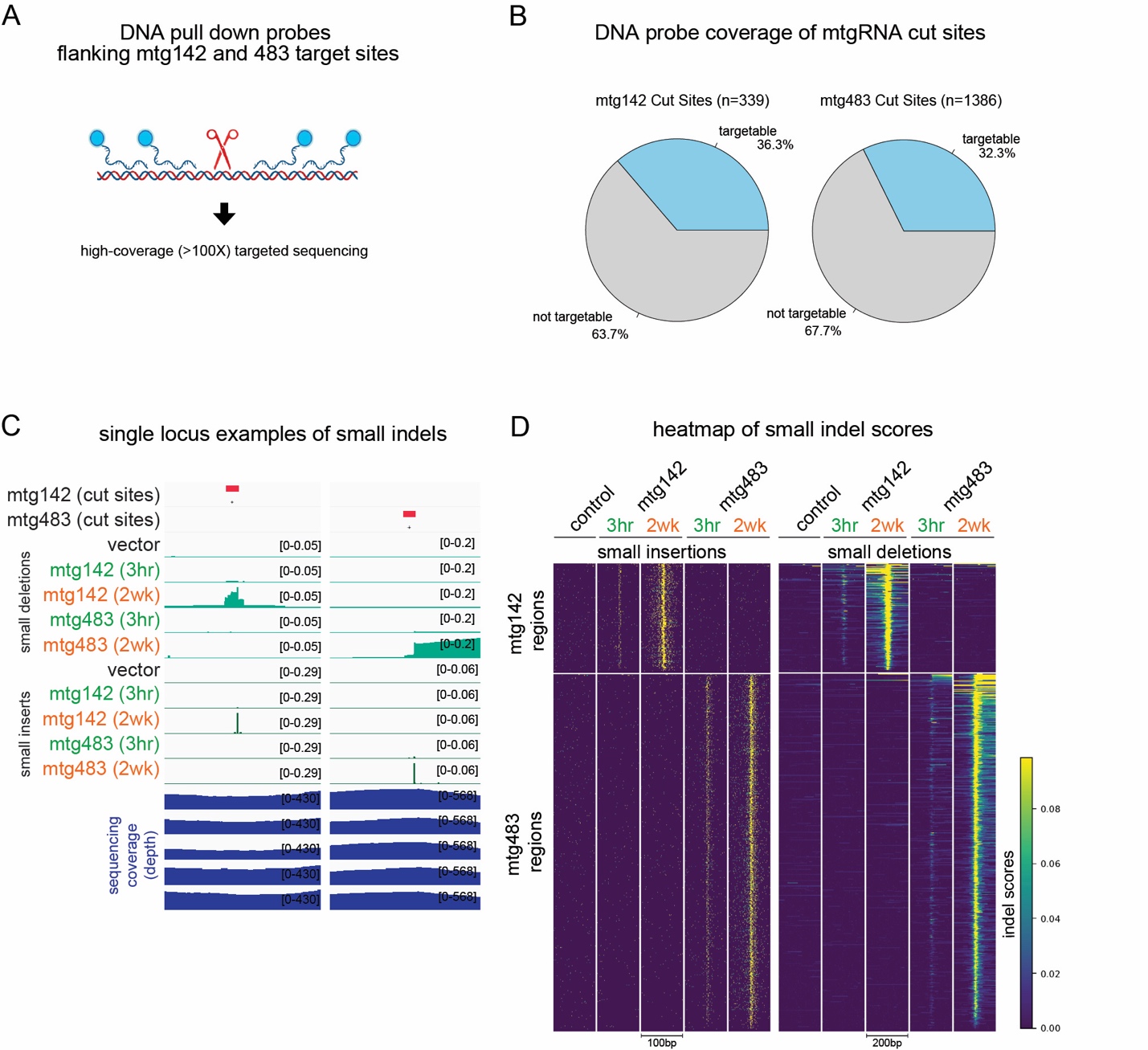

Fig. S10. Supplementary data for Figure 4 showing DNA enrichment strategy for deep sequencing of cut sites and small indel detection

A) Schema of DNA probe library design: multiple DNA-based and affinity tagged probes were designed to flank cut sites. After pull-down, DNA was subject to high coverage sequencing (>100X).
B) Pie charts summarizing the number of cut sites that were targetable using the strategy outlined in **(A)**. Due to the degeneracy of similar cut sites, ~36% and ~32% of all mtg142 and mtg483 sites could be covered using this strategy while maintaining target site specificity.
C) Two single locus examples of mtgRNA cut sites showing small deletions or insertions that develop by 2 weeks at the target sites. Coverage of the two regions show ~400-500X coverage.
D) Heatmap of small insertions and deletions at mtg142 and mtg483 regions.

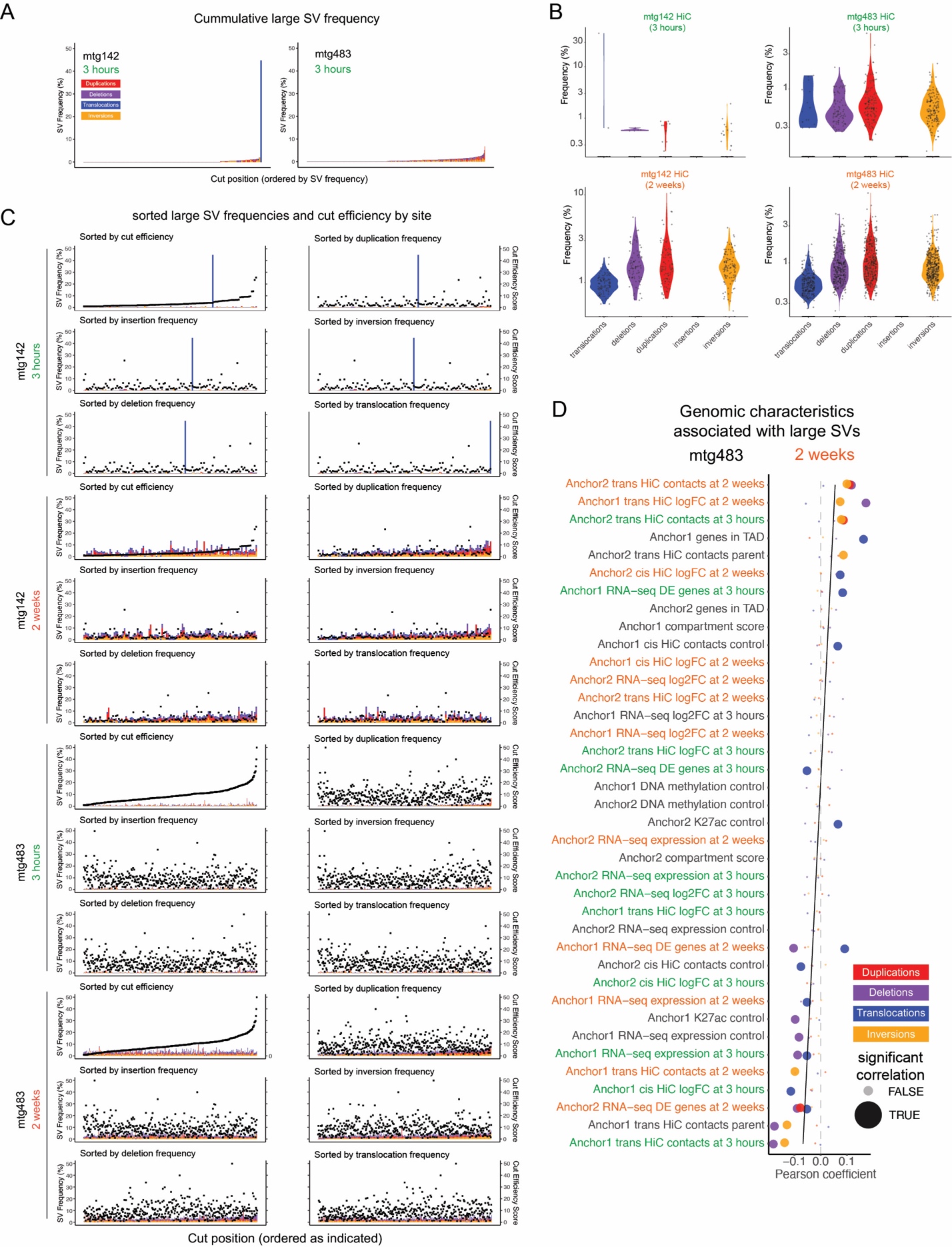

Fig. S11. Supplementary data for Figure 4 showing additional timepoints, SV frequency comparisons, and mtg483 SV frequency correlation with genomic alterations.

A) Cumulative large SV frequency (y-axis) ordered by total frequency along the x-axis. Each stacked bar represents a cut site. The three hour time points are shown.
B) Large SV frequency (left y-axis) ordered by cut efficiency or frequency of individual SV types along the x-axis. The right y-axis shows the cut efficiency score. Each stacked column represents a cut site.
C) Violin plots of large SV frequency by condition and timepoint showing overall SV frequencies.
D) Correlation coefficients of genomic measurements at control (baseline), 3 hours and 2 weeks with SVs that are detected at 2 weeks in condition mtg483. Anchor 1 and 2 demark either end of the SV breakpoints (ordered numerically by chromosome number and location). Anchor specific genomic measurements (y-axis) are ordered by average Pearson correlation coefficient of the 4 indicated large SVs (x-axis).

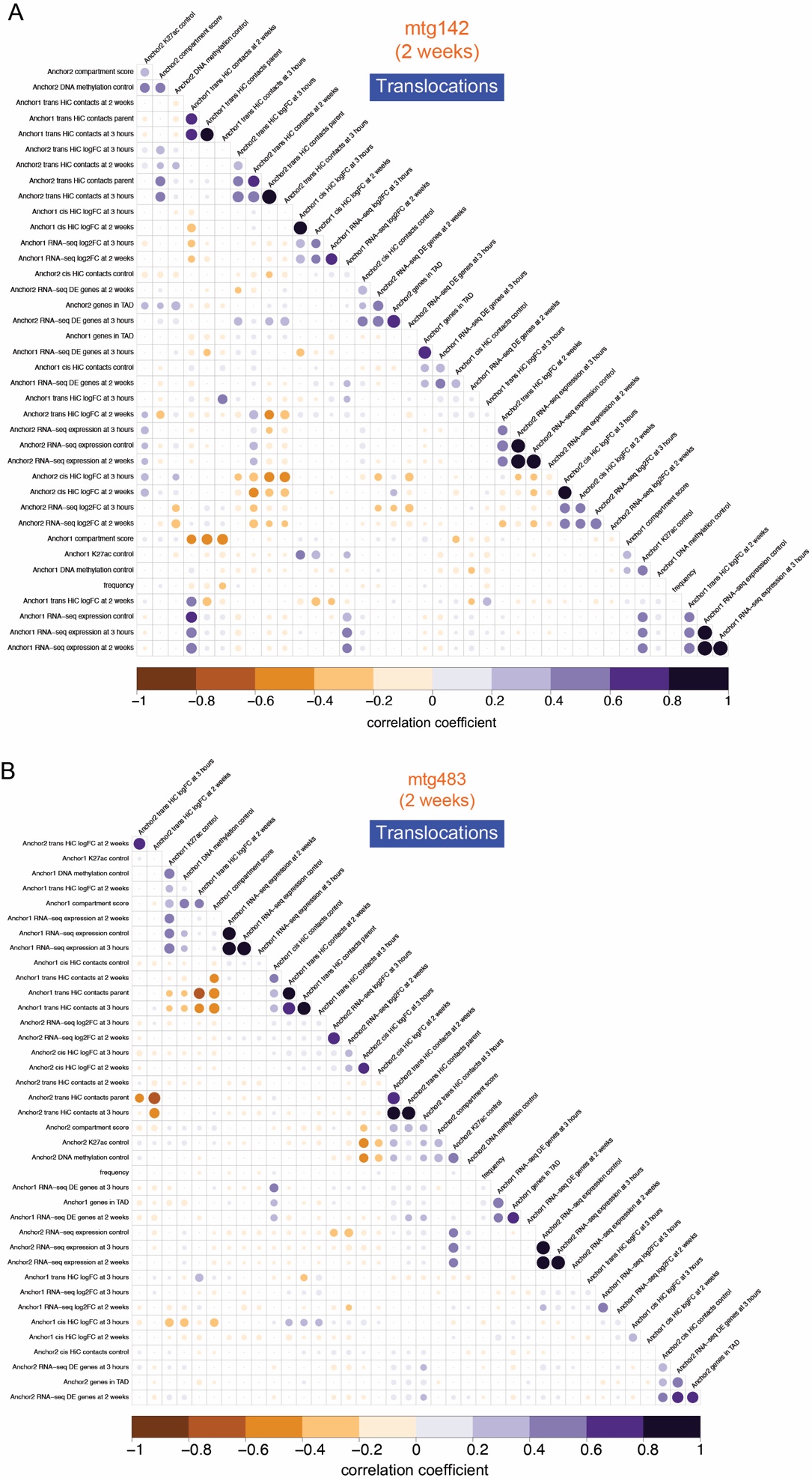

Fig. S12. Supplementary data for Figure 4C showing all genomic correlations and translocation Frequency

A) and **B)** Correlation plot for mtg142 and mtg 483 at two weeks subset by translocations. Dots quantify the correlation coefficient of pairwise comparisons between genomic measurements. Dot size correlates with coefficient magnitude and color with direction. The absence of a dot indicates the correlation was non-significant (significance threshold p < 0.05 using Student's t-distribution transformation of correlation coefficients). Features were hierarchically clustered using complete linkage clustering to identify patterns of related correlations.

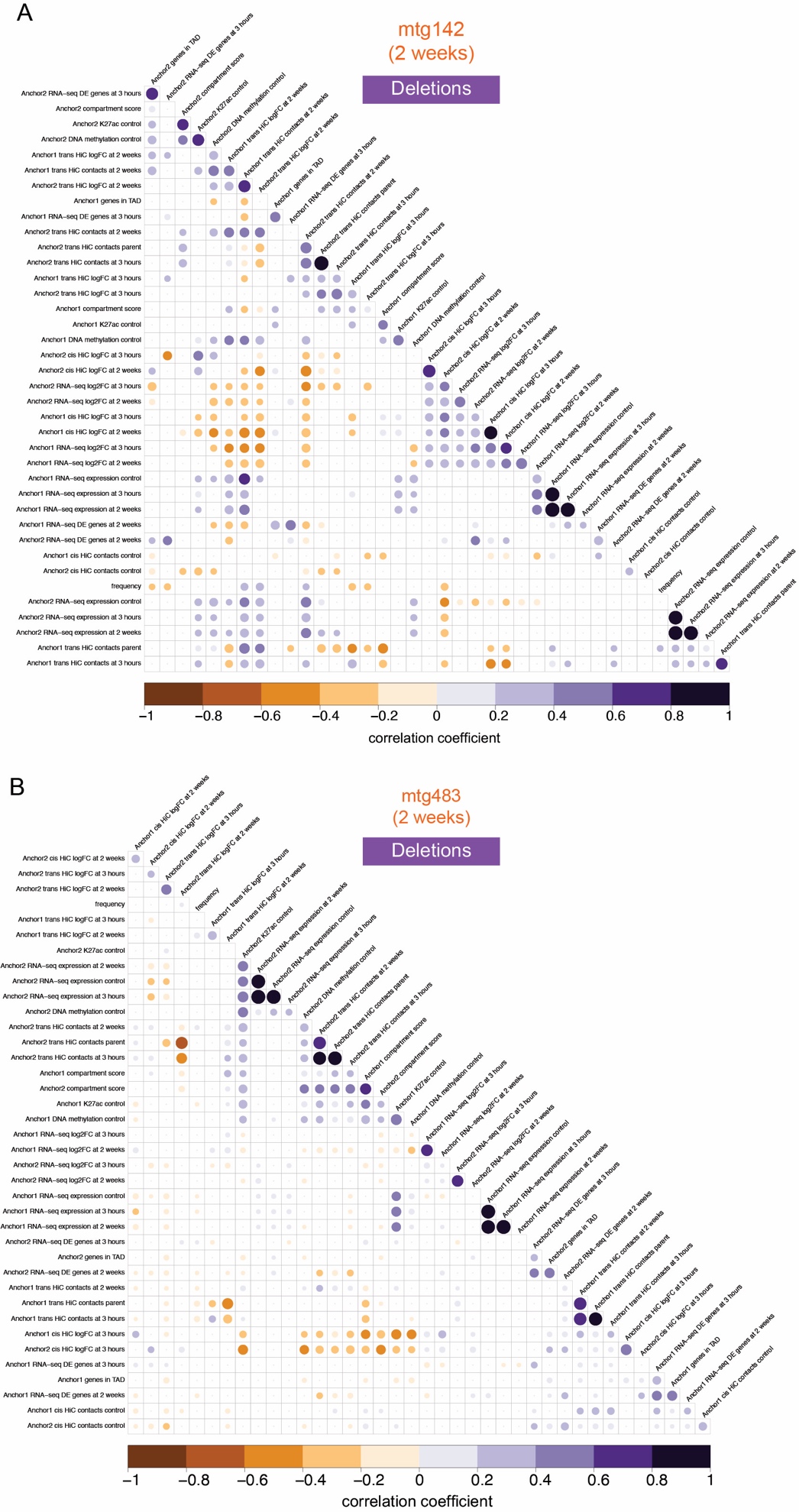

Fig. S13. Supplementary data for Figure 4C showing all genomic correlations and deletion Frequency

A) and **B)** Correlation plot for mtg142 and mtg 483 at two weeks subset by deletions. Dots quantify the correlation coefficient of pairwise comparisons between genomic measurements. Dot size correlates with coefficient magnitude and color with direction. The absence of a dot indicates the correlation was non-significant (significance threshold p < 0.05 using Student's t-distribution transformation of correlation coefficients). Features were hierarchically clustered using complete linkage clustering to identify patterns of related correlations.

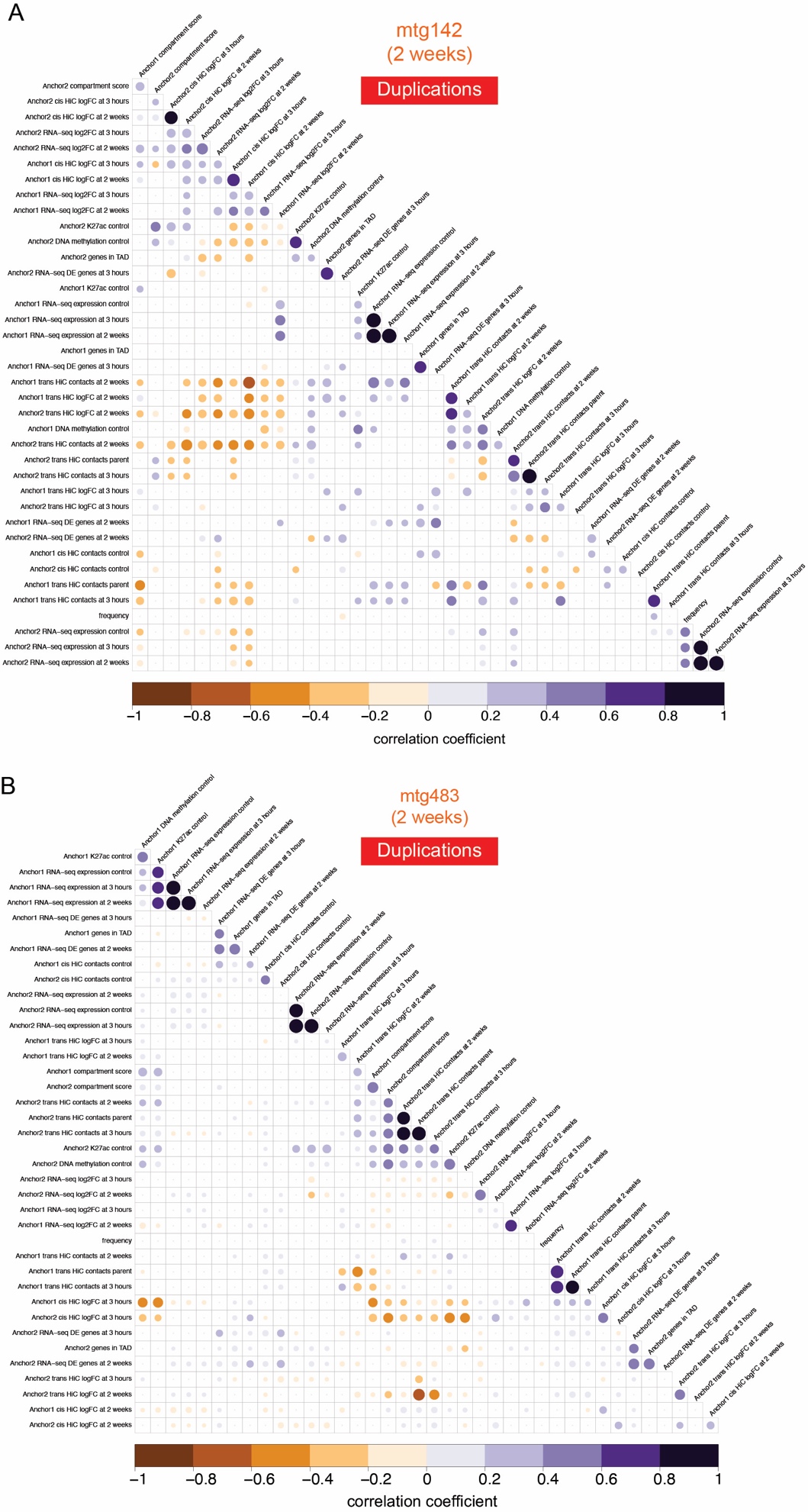

Fig. S14. Supplementary data for Figure 4C showing all genomic correlations and duplication Frequency

A) and **B)** Correlation plot for mtg142 and mtg 483 at two weeks subset by duplications. Dots quantify the correlation coefficient of pairwise comparisons between genomic measurements. Dot size correlates with coefficient magnitude and color with direction. The absence of a dot indicates the correlation was non-significant (significance threshold p < 0.05 using Student's t-distribution transformation of correlation coefficients). Features were hierarchically clustered using complete linkage clustering to identify patterns of related correlations.

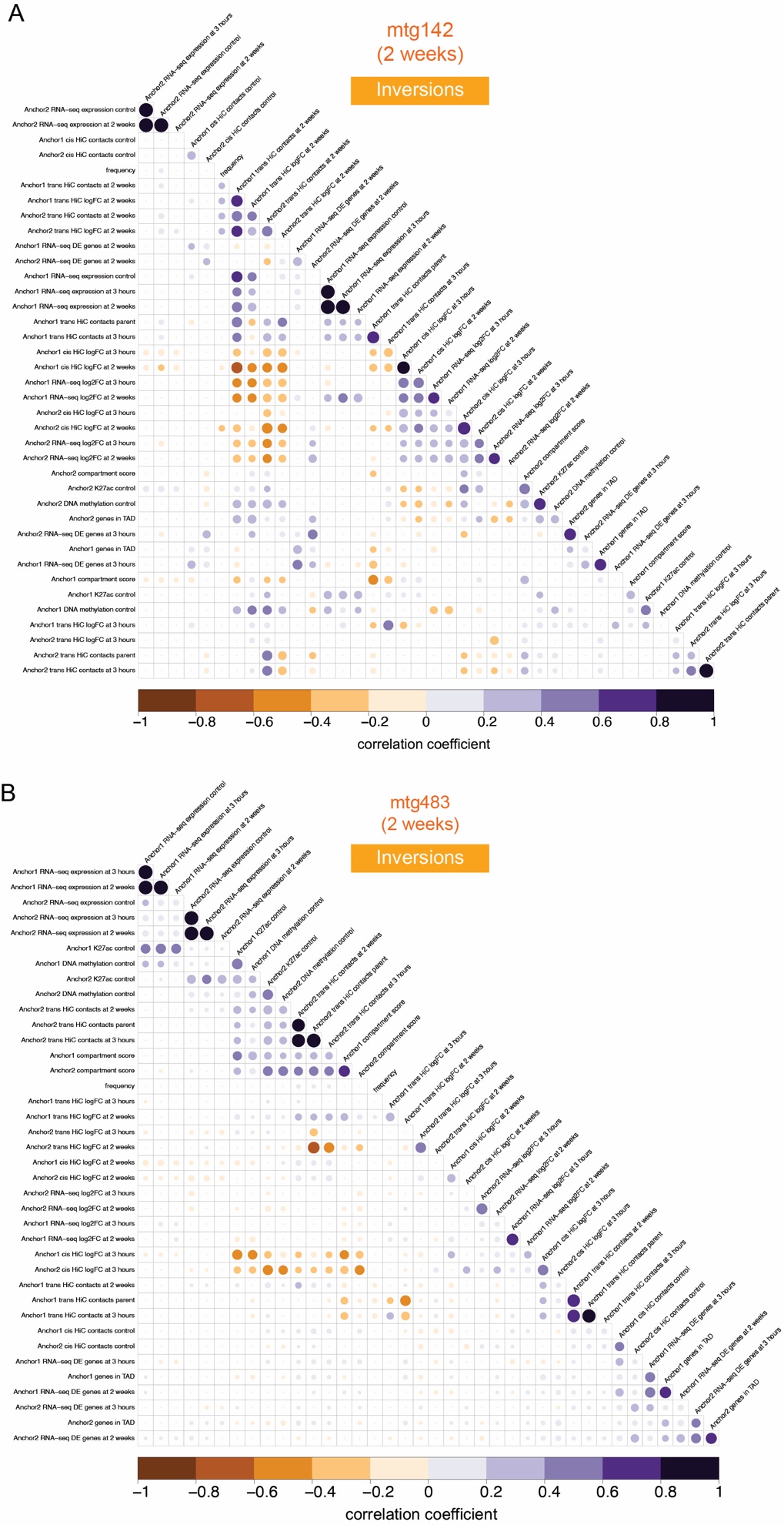

Fig. S15. Supplementary data for Figure 4C showing all genomic correlations and Inversion Frequency

A) and **B)** Correlation plot for mtg142 and mtg 483 at two weeks subset by inversions. Dots quantify the correlation coefficient of pairwise comparisons between genomic measurements. Dot size correlates with coefficient magnitude and color with direction. The absence of a dot indicates the correlation was non-significant (significance threshold p < 0.05 using Student's t-distribution transformation of correlation coefficients). Features were hierarchically clustered using complete linkage clustering to identify patterns of related correlations.

Data S1 to 6 (separate file)

Data Tables S1 to S4

Gene-set enrichment analysis of mtg142 and mtg483 at all timepoints

Data Tables S5 to 6

DNA probes used for pull-down sequencing of mtg142 and mtg483 cut sites
